# Regulation of thymocyte β-selection, development and positive selection by glycogen synthase kinase-3

**DOI:** 10.1101/712521

**Authors:** Michael J. Parsons, Satish Patel, Bradley W. Doble, Pamela S. Ohashi, James R. Woodgett

## Abstract

Glycogen synthase kinase-3 (GSK-3) is a ubiquitously expressed serine/threonine kinase, that exists as two isoforms in mammals, GSK-3α and GSK-3β, that are key downstream mediators of the phosphatidylinositol 3’ kinase, Wnt, Notch and other pathways. Here, we report that simultaneous inactivation of both GSK-3α and GSK-3β during early thymocyte ontogeny has profound effects on both β-selection and positive selection, key checkpoints essential to producing functionally mature αβ T cells. Conditional GSK-3α/β knockout animals (LckCre^+^ GSK-3αβ^fl/fl^) possessed pre-double positive (pre-DP) thymocytes (CD4^−^CD8^−^CD117^−^CD25^−^) with compromised TCRβ chain expression along with elevated levels of β-catenin and reduced Notch activity. β-selection was impaired allowing pre-DP thymocytes to differentiate to DP thymocytes (CD4^+^CD8^+^) while bypassing strict requirements for productive TCRβ chain rearrangements and functional expression. Also impaired was the requisite pre-TCR and Notch-mediated expansion that normally precedes differentiation to the DP stage. Consequently, LckCre^+^ GSK-3αβ^fl/fl^ mice initially generated fewer DP thymocytes that expressed significantly reduced levels of mature TCR. The aberrant DP thymocytes expressed high levels of the pro-survival Bcl-2 family member Mcl-1, failed positive selection and accumulated as CD4^hi^CD8^lo^ positive selection intermediates resulting in loss of both mature CD4 and CD8 lineages. LckCre^+^ GSK-3αβ^fl/fl^ mice succumbed to oligoclonal peripheral lymphomas with high penetrance. These data reveal essential roles for GSK-3 in several checkpoints of early T cell development.

## Introduction

Thymocyte development requires navigation of cells through a well-orchestrated series of differentiation and maturity events that are modulated by multiple signalling pathways and results in the selection of a functional, but not auto-reactive, T cell repertoire, *ab initio*. Early stages of αβ thymocyte development prior to CD4 and CD8 expression (CD4^−^CD8^−^ (double-negative, DN) sub-population), have been separated into four stages (DN1-DN4) based on expression of CD117, CD44 and CD25. A critical checkpoint known as β-selection occurs at the DN3 stage, phenotypically defined as being CD4^−^ CD8^−^ CD117^−^ CD44^−^ CD25^+^ ^1–3^. It is at this stage that rearrangements within the TCRβ chain locus produce a diverse repertoire of TCRβ chains with varied specificities including non-functional variants. Productively rearranged TCRβ chains go on to form complexes with invariant pre-Tα chains and CD3/ζ/η components to form what is referred to as the pre-TCR^4–6^. Signalling from the pre-TCR promotes survival of thymocytes that would otherwise die by neglect, along with down-regulation of CD25 to produce DN4 (CD4^−^ CD8^−^ CD117^−^ CD44^−^ CD25^−^) thymocytes. Pre-TCR signalling also triggers a series of other events critical for transition to double-positive (DP) cells expressing both CD4 and CD8 that includes TCRβ chain allelic exclusion, differentiation and proliferation. Only thymocytes that have undergone successful rearrangement at the TCRβ locus receive the appropriate proliferative, survival and differentiation signals necessary to transition from CD117^−^ CD44^−^ CD25^−^ CD4^−^CD8^−^ TCRβ^+^ (pre-DP) to CD4^+^CD8^+^ (DP) thymocytes^7, 8^. Numerous studies have shown that various mouse models including: RAG1^null^, RAG2^null^, TCRβ^−/−^ and pTα^−/−^ mice are unable to form the pre-TCR and consequently arrest development at the DN3 stage^9–12^.

Several signalling pathways including Notch, CXCR4, p53, NFAT and Egr-3 have been shown to play important roles during β-selection^13–22^. Specifically, one model of Notch function posits that this molecule works in delicate balance with the pre-TCR by providing signals associated with the proliferative-self renewal of early post β-selected thymocytes prior to pre-TCR directed differentiation. Notch activity is transient and eventually gives way to signals from the pre-TCR that promote differentiation involving loss of CD25 and pTα expression along with cell cycle arrest, RAG activation in preparation of mature TCRα gene rearrangement and, finally, up-regulation of CD4 and CD8^2, 3, 23^.

The phosphatidylinositol 3’ kinase (PI3K) pathway also plays a key role during thymocyte development. Downstream of PI3K are AGC kinases such as phosphoinositide-dependent kinase 1 (PDK1) and protein kinase B (Akt/PKB)^24^. Studies targeting each of these components of PI3K signalling have revealed important contributions to pre-TCR and mature TCR signalling^25–28^. GSK-3 activity is negatively regulated through Akt-mediated phosphorylation at serines 21 and 9 (GSK-3α and β respectively), thus linking GSK-3 regulation to signals acting through PI3K^29^.

While it has been reported that constitutive activation of Akt in absence of pre-TCR and NOTCH signals can prevent programmed cell death of DN thymocytes independent of several Bcl-2 members there is evidence that Bcl-2 family members such as Mcl-1 play a crucial role in T cell survival, development and tumorigenesis^30–35^. GSK-3 regulates Mcl-1 stability through ubiquitin-mediated degradation^34, 35^. Deletion of Mcl-1 impairs thymocyte survival causing reduced DP cell number and elevated apoptosis for DN and DP subsets. Only a small fraction of DP thymocytes receive the appropriate amount of TCR signal and are selected for further differentiation into mature CD4 and CD8 lineages in a process referred to as positive selection^36–38^.

In mammals, GSK-3 exists as two isoforms termed GSK-3α (51kDa) and GSK-3β (47kDa) encoded by distinct genes that are implicated in multiple pathways and diseases such as diabetes, Alzheimer’s disease, bipolar disorder and cancer^39, 40^. GSK-3β^null^ embryos die late in development while GSK-3α^null^ mice are viable though males are sterile^41–43^. Regulation of GSK-3 activity is atypical as it is negatively regulated in response to cellular signals leading to inactivation of GSK-3 and de-repression of certain substrates^44, 45^.

Wnt signalling regulates GSK-3 through a distinct mechanism and many studies employing *gain-of-function* and *loss-of-function* mutations have demonstrated that Wnt-signalling and its central effector, β-catenin, figures prominently in developmental processes including thymopoesis^46–48^. In the thymus Wnt proteins, secreted mainly by the thymic epithelium, signal through the Frizzled/LRP5/6 co-receptor complex to divert GSK-3 activity to LRP5/6, allowing β-catenin to accumulate and associate with TCF/LEF transcription factors^49, 50^. TCF1/LEF1^null^ mice have impaired thymocyte development while ICAT, a natural inhibitor of β-catenin and TCF interaction, has been shown to block the DN to DP transition^51–53^. Signals from the pre-TCR and TCR lead to β-catenin stabilization and activation^54–60^.

Given the association of GSK-3 with signalling pathways shown to play key roles during thymocyte development, we crossed floxed alleles of GSK-3α and GSK-3β with transgenic animals expressing Cre-recombinase under control of the p56^lck^ proximal promoter^61^ to selectively inactivate the GSK-3 isoforms early in T cell development. While inactivation of one isoform or the other has little effect, loss of both GSK-3α and GSK-3β isoforms disrupted key regulators of thymocyte β-selection. We show that pre-DP’s differentiate to form aberrant DP’s that lack mature TCR without the normally requisite pre-TCR associated expansion and result in almost complete loss of both CD4 and CD8 mature T cell lineages.

## Materials and Methods

### Mice

Creation of GSK-3α^flox/flox^ and GSK-3β^flox/flox^ mice was previously described and were bred together to generate various allelic combinations of floxed GSK-3α and β isoforms^43, 44, 49^. GSK-3α^fl/fl^β^fl/fl^ (LckCre^+^ GSK-3αβ^fl/fl^) mice were subsequently bred to additional strains including transgenic p56LckCre, transgenic TCR LCMV-P14 and RAG1^null^ ^10, 61–65^.

### Flow cytometry and antibodies

The following antibodies were obtained from either ebioscience or BD Bioscience and used for surface staining: CD117 PerCP-efluor 710 (clone2B8), CD4FITC (GK1.5), CD4APC (clone RM4-5), CD25 APC (clone 3C7), CD24PE and CD24PE-CY5, (clone M1/69), CD3APC (clone 145-2C11), CD69PE (clone H1.2F3), CD5 FITC (clone53-7.3), CD71PE (clone C2), TCRβPE (clone H57-597), TCRγδ FITC (clone UC7-13D5), TCRvα-2 (clone B20.1), TCRvβ8.1/8.2 (clone F23.1), pTα biotin (BD cat # 552407) (conjugated with streptavidin PE). Experiments focused on CD4^−^CD8^−^(DN) populations used a lineage exclusion (lin^neg^) panel of antibodies conjugated with either FITC or PE: (CD4, CD8, ter119, TCRγδ, CD11b, DX5 (pan NK), Gr1, CD19).

Cells were first blocked using blocking antibody CD116/32 (ebiosciences cat # 14-0161-86) and then stained with saturating concentrations of titrated antibody. Data was acquired in FCS 3.0 file format on a Beckman Coulter Gallios analyser except experiments analysing apoptosis using Annexin V that were performed on a BD Biosciences LSRII. In general antibody capture beads (BD Biosciences cat # 552845 and 552843) were used with bound antibody as compensation controls except for cell viability dye where cells were used. Live cells were gated according to their forward-scatter and side scatter profiles along with exclusion based on the viability dye SYTOX-blue (Invitrogen cat# S34857) where possible. Doublets were removed though forward scatter time of flight (width) versus peak (height) profile. Fluorescence minus-one controls were used to set gates as necessary. Analysis was performed using Beckman Coulter Kaluza analysis software except where indicated. Representative bivariate plots and histograms are shown with figure legends indicating associated N numbers, means with standard deviations and relevant “P” values (nonparametric one tailed) denoting significance.

### Intracellular staining

Intracellular staining for TCRβ involved blocking with CD116/32 blocking antibody along with purified TCRβ to ensure no surface TCRβ was stained followed by staining for CD117 and CD25 in addition to the lineage exclusion panel. After surface staining cells were washed with stain buffer, treated with cytofix-cytoperm (BD) for 20 min then washed twice with perm-wash buffer (BD). Permeabilized cells were incubated with either TCRβ (BD) or β-catenin FITC antibodies (BD transductionl aboratories cat # 610156) for 45min in perm-wash buffer. Finally, cells were washed twice in perm-wash buffer and re-suspended in stain buffer for analysis.

### Genomic PCR

Thymocytes from LckCre^+^GSK-3α^fl/fl^ β^fl/fl^ and LckCre^−^GSK-3α^fl/fl^ β^fl/fl^ were sorted into DN3, DN4, DP, CD4 (SP), and CD8 (SP) sub-populations. 1X10^5^ cells were lysed in 50μl of a buffer containing 1X PCR buffer, 0.5% NP40, 0.5% Tween 20 and Proteinase K (10mg/ml). PCR reactions were performed using the following primer sets: GSK-3α forward (cccccaccaagtgatttcactgcta) and reverse (aacatgaaattccgggctccaactctat), GSK-3β forward (ggggcaaccttaatttcatt) and reverse (tctgggctatagctatctagtaacg). Larger bands are either the GSK-3α or GSK-3β floxed alleles while the smaller bands are the excised exon 2 fragment from either the GSK-3α or GSK-3β floxed alleles.

### Western blotting

Thymocytes from LckCre^+^GSK-3α^fl/wt^ β^fl/fl^ (β^3/4^), LckCre^+^GSK-3α^fl/fl^ β^fl/wt^ (α^3/4^), LckCre^−^GSK-3α^fl/fl^ β^fl/fl^ and LckCre^+^GSK-3α^fl/fl^ β^fl/fl^ mice were sorted for the CD4^+^CD8^+^ (DP) thymocyte subset. These cells were then lysed in T-PER tissue protein extraction reagent (Pierce) containing protease inhibitor cocktail (Roche) and phosphatase inhibitor cocktail 1(Sigma), run on a 10%SDS PAGE gel and probed for GSK-3αβ(1:10,000 Biosource), β-catenin (1:1,000 BD transduction labs), Mcl-1 (1:500 Santa Cruz) and GAPDH (1:100,000 Abcam)

### CFSE proliferation assay

Anti-CD3 and anti-CD28 antibodies were bound to 96 well MaxiSorp plates (Nunc) overnight at 4°C using concentrations of 10μg/ml and 5μg/ml respectively. The plates were then washed in PBS and blocked in RPMI 1640 media containing 10% low immunoglobulin serum, L-glutamine (2mM), and 2-mercaptoethanol (50μM). IL-2 was added at a concentration of 0.8ng/ml, as required. Spleen cells with lysed red blood cells were incubated with CFSE at a concentration of 10μM for 15 min at 37°C then washed with media before plating at 1 × 10^5^ cells in 100μl of media in each well. Cells were cultured for 72 hrs and then stained for CD4 and CD8. Untreated cells were also stained for CD4, CD8, CD3 and TCRβ to determine the status of CD3 and TCRβ prior to stimulation.

### TCRVβ repertoire clone screen

Spleen cells from 5 month animals were sorted for CD8 positive cells and cDNA was prepared using SuperScript III First-Strand Synthesis System for RT-PCR from Invitrogen (cat # 18080-051). The cDNA was then used in the TCR*Express*™ MouseTCRVβ screening kit (BioMed Immunotech) using procedures provided. Additional TCRVβ analysis was carried out using the mouse Vβ TCR screening panel kit provided by BD Biosciences (cat # 557004).

### Thymocyte survival assay

To assess spontaneous death 1X10^6^ thymocytes were plated in media for 24, 48 and 72 hrs. Thymocytes were then washed, blocked and stained for CD4 and CD8 to identify DP’s. Apoptosis was assessed using the Annexin V-FITC apoptosis detection kit (BD cat # 556570). For anti-CD3-induced death thymocytes were plated at 0.5 × 10^6^ cells/well of a 96 well plate (Nunc Maxisorb) coated with anti-CD3 antibodies (clone 145-2C11) and cultured for 24 and 48 hrs. For anti-Fas-induced death cells were plated similarly except in the presence of anti-Fas (2μg/ml) (BD cat # 554254) and protein G (2μg/ml) for 30 min. For dexamethasone-induced death, 2 × 10^6^ thymocytes were cultured for 3 hrs with 4ml of media containing dexamethasone at various concentrations (1.0μM, 0.1μM, 0.01μM or 0μM). Following these treatments cells were washed, blocked and stained for CD4 and CD8 to identify DP and DN cells. Apoptosis was also assessed using the Annexin V-FITC apoptosis detection kit (BD cat # 556570).

### Real Time PCR

Mouse CD4 Dynabeads (Invitrogen cat# 114.45D) were used to enrich the CD4^−^CD8^−^ (DN) fraction of thymocytes from 3-4 wk old mice which were then sorted for the CD117^−^ CD25^−^ lin^neg^ (DN4) subpopulation. DP thymocytes were sorted according to CD4 and CD8 co-expression. 7AAD was included in sorts to ensure viability. RNA was extracted from CD117^−^ CD25^−^ lin^neg^ (DN4) and CD4^+^CD8^+^ (DP) thymocytes using the RNeasy Plus Mini Kit (Qiagen) and SuperScript lll reverse transcriptase was used to make cDNA using the protocol supplied by Invitrogen (cat # 18080-093). Power SYBR Green PCR master mix (Applied Biosystems cat # 4367659) was used to prepare samples before running on an Applied Biosystems 7500 Real-Time PCR System. Analysis was performed using the comparative C_T_ method (ΔΔC_T_ Method) described by Applied Biosystems with GAPDH used as the endogenous reference gene. PCR product values were expressed relative to LckCre^−^ GSK-3 control samples. An average of three wells was used to calculate PCR product values for each experiment and each experiment was repeated at least four times with standard error used to measure variability.

Primer sets used:

EGR3 fwd (ccggtgaccatgagcagttt) rev (taatgggctaccgagtcgct);
Bcl-X_L_ fwd (ggatctctacgggaacaatctt) rev (gtcaggaaccagcggttga);
HES1 fwd (ccagccagtgtcaacacga) rev (aatgccgggagctatctttct);
DTX1 fwd (cctacatcatcgacctacaatcc) rev (ccacacgatgcccttgcccg)
GAPDH fwd (tgaccacagtccatgccatc) rev (gacggacacattgggggtag)

For TCRβ rearrangements the following primer sets were used:

Dβ2(gtaggcacctgtggggaagaact)-Jβ2(tgagagctgtctcctactatcgatt)
Vβ5.1-Jβ2(gtccaacagtttgatgactatcac)-Jβ2(tgagagctgtctcctactatcgatt)
Vβ8.2(cctcattctggagttggctaccc)-Jβ2(tgagagctgtctcctactatcgatt)

Supplemental supporting data are available on-line (see Supplementary Data).

## Results

### Generation of mice harbouring early thymocyte-specific deletion of GSK-3α and GSK-3β

Floxed GSK-3α and GSK-3β mice were crossed to transgenic animals expressing Cre-recombinase under control of the p56^lck^ proximal promoter. Inter-crosses between floxed LckCreGSK-3α and floxed LckCreGSK-3β mice produced T-cell conditional mice with disrupted GSK-3 expression according to various allelic combinations of GSK-3α and GSK-3β to allow investigation of potential isoform-specific differences and/or gene dosage effects on thymocyte development (**Suppl. Fig. 1**).

Thymocytes from LckCre^+^ GSK-3αβ^fl/fl^ and littermate LckCre^−^ GSK-3αβ^fl/fl^ mice were sorted into CD117^−^ CD25^+^ lin^neg^ (DN3), CD117^−^ CD25^−^ lin^neg^ (DN4), CD4^+^CD8^+^ (DP), CD4^+^CD8^−^ (CD4 SP) and CD4^−^CD8^+^ (CD8 SP) subsets and genomic PCR was performed to ascertain the extent of either GSK-3α or GSK-3β exon 2 excision. Significant excision of exon 2 of GSK-3α and GSK-3β was evident by the DN3 stage of thymocyte development for LckCre^+^ GSK-3αβ^fl/fl^ mice. This is in keeping with the well-documented activity of the p56^lck^ proximal promoter, which reaches maximal expression around the DN3 stage of thymocyte development^61, 62, 65^. Immunoblots of DP thymocytes from LckCre^+^ GSK-3α^fl/fl^ β^fl/wt^ (α^3/4^), LckCre^+^ GSK-3α^fl/wt^ β^fl/fl^ (β^3/4^), LckCre^+^ GSK-3αβ^fl/fl^ and LckCre^−^ GSK-3β^fl/fl^ mice were probed for both isoforms of GSK-3, confirming isoform-selective loss of appropriate GSK-3 proteins (**Suppl. Figs. 2a, 2b**).

### Inactivation of both GSK-3α and GSK-3β causes marked loss of mature T cells, reduced thymocyte cell number and accumulation of positive selection intermediates

Comparison between LckCre^+^ GSK-3αβ^fl/fl^ and littermate LckCre^−^ GSK-3αβ^fl/fl^ control mice at 3 wks of age showed significant loss of mature T cells in mice having lost both isoforms of GSK-3 when first analyzed in terms of CD4 and CD8 surface expression in the thymus. We also noted an increase in the proportion of DN cells (13.2% vs. 3.5%) indicating a potential block in development prior to DP differentiation. Subsequent analysis of the CD3^+^CD24^−^ subpopulation, which identifies mature thymocytes revealed the extent to which mature CD4 and CD8 (SP) thymocytes are missing from the LckCre^+^ GSK-3αβ^fl/fl^ thymus (CD4: 3.7% vs. 61.8% and CD8: 5.4% vs. 34.7%). Furthermore, the majority of CD3^+^CD24^−^ thymocytes from LckCre^+^ GSK-3αβ^fl/fl^ mice continue to express both CD4 and CD8, which is another indication of aberrant development. Analysis of mice at 6-8 wks of age showed similar findings (**Fig. 1a, 1b, 1d and Suppl. Fig. 2c**). Thymocyte-specific loss of either GSK-3α or GSK-3β alone had no discernible effect on thymocyte development as defined through CD4 vs. CD8 expression. Notably even the presence of a single allele of either GSK-3α (GSK-3α^fl/fl^ β^fl/wt^ (α^3/4^)) or GSK-3β (GSK-3α^fl/wt^ β^fl/fl^ (β^3/4^)) in the presence of the p56^lck^Cre transgene produced mice with entirely normal ratios and cell numbers of DN, DP, CD4 (SP), CD8 (SP) sub-populations along with normal levels of CD3 and TCR expression in the thymus, compared to littermate controls. These data show functional redundancy between GSK-3α and GSK-3β with a single allele of either GSK-3α or GSK-3β sufficient to mediate the complex signalling outcomes governing thymocyte development. Given the severity of the phenotype observed for LckCre^+^ GSK-3αβ^fl/fl^ mice versus the lack of obvious phenotype for either of the singly deleted animals, we focused further characterization on the LckCre^+^ GSK-3 αβ^fl/fl^ mice (**Fig. 1c, Suppl. Fig. 3a and data not shown**).

**Figure 1.**
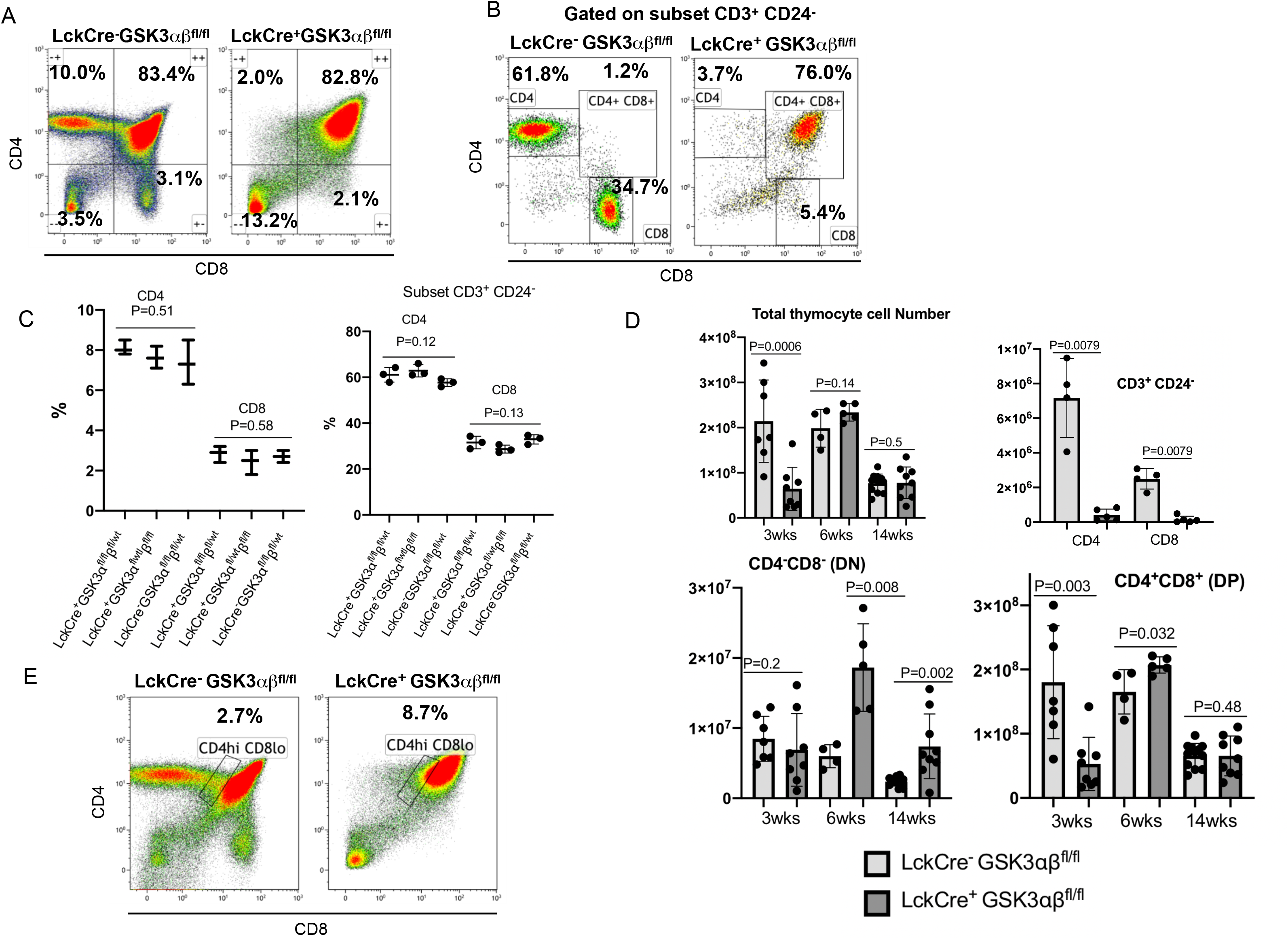
Loss of GSK-3 leads to a significant reduction in mature T-cell production and overall thymocyte cellularity. (a,b) (N=10) LckCre^−^ GSK-3αβ^fl/fl^ and (N=11) LckCre^+^ GSK-3αβ^fl/fl^ thymocytes of 3 week old mice were stained for CD4 vs. CD8 expression along with a CD4 vs. CD8 stain of mature T cell sub-populations gated on the CD3^+^/CD24^−^ subset. Quadrant statistics indicate proportion of cells in each quadrant (a. mean DN 3.3% +/− 0.74 vs. 11 +/− 5.3 P<0.0001, mean DP 82.4% +/− 7.2 vs. 82% +/− 7.0 P =0.458 : b. CD3^+^ CD24^−^ Gate, CD4 mean 54.9% +/− 10.3 vs. 3.0% +/− 1.8 P =0.0007, CD8 mean 28% +/− 3.7 vs. 3% +/− 2.1 P=0.0002, DP mean 2.8% +/− 1.1 vs. 48% +/− 24 P=0.0002) (c) (N=3) LckCre^+^ GSK-3α^fl/fl^β^fl/wt^, (N=3) LckCre^+^ GSK-3α^fl/wt^β^fl/fl^ and (N=3) LckCre^−^ GSK-3α^fl/fl^β^fl/wt^ thymocytes of 3 week old mice were stained for CD4 vs. CD8 expression along with a CD4 vs. CD8 stain of mature T cell sub-populations gated on the CD3^+^/CD24^−^ subset. Histogram comparisons show no significant difference between populations (CD4^+^ P=0.51, CD8^+^ P=0.58, CD4^+^CD3^+^CD24^−^ P= 0.12 and CD8^+^CD3^+^CD24^−^ P=0.13. (d) Comparison of total, DP, and DN thymocyte numbers for (N=10) LckCre^−^ GSK-3αβ^fl/fl^ and (N=11) LckCre^+^ GSK-3αβ^fl/fl^ mice at 3, 6-7 and 14 weeks of age (total 3 week P= 0.0006, 6-7 week P=0.14 and 14 week P=0.5 respectively, DN 3 week P=0.2, 6-7 week P=0.008, 14 week P=0.002, DP 3 week P=0.003, 6-7 week P=0.032, 14 week P=0.48). Error bars given as standard error. Comparison of thymocyte numbers for CD4^+^/CD3^+^/CD24^−^ and CD8^+^/CD3^+^/CD24^−^ SP thymocytes at 6-7 weeks of age (P= 0.0079 and P=0.0079 respectively). Error bars given as standard error. (e) Increase in proportion of (N=11) LckCre^+^ GSK-3αβ^fl/fl^ CD4^hi^/CD8^lo^ thymocytes compared to (N=10) LckCre^−^ GSK-3αβ^fl/fl^ thymocytes at 3 weeks of age. Numbers in plot represent percent CD4^hi^/CD8^lo^ cells (mean 6.7% +/− 4.1 vs. 2.5% +/− 1.3 P=0.0027).

Examination of total thymic cell number revealed a ~3-fold loss of cellularity in 3 wk old LckCre^+^ GSK-3αβ^fl/fl^ mice compared to littermate LckCre^−^ GSK-3αβ^fl/fl^ mice. This was due to an absolute loss of DP and SP thymocytes since the number of thymocytes in the 3 wk old DN compartment did not change relative to LckCre^−^ GSK-3αβ^fl/fl^ mice. Further comparison of DP and DN cell numbers between 3, 6 and 14 wk old animals revealed a striking increase in DP cellularity in LckCre^+^ GSK-3αβ^fl/fl^ mice at six weeks of age, in addition to an increase in DN cells. However, these animals remained unable to produce mature T cells as judged by comparison of CD4^+^ and CD8^+^ thymocyte numbers within the CD3^+^CD24^−^ sub-population. LckCre^+^ GSK-3αβ^fl/fl^ DP cell numbers did not continue to rise unchecked. There was a return to lower LckCre^+^ GSK-3αβ^fl/fl^ DP cell numbers, comparable to LckCre^−^ GSK-3αβ^fl/fl^ animals by 14 wks of age, suggesting that processes governing age-associated thymic atrophy remained intact (**Fig. 1d**)^67^. We also observed an increase in the fraction of CD4^hi^CD8^lo^ cells in LckCre^+^ GSK-3αβ^fl/fl^ mice at 3 wks of age (8.7% vs. 2.7%) suggesting that thymocytes of LckCre^+^ GSK-3αβ^fl/fl^ mice are unable to complete positive selection, resulting in accumulation of positive selection intermediates (**Fig. 1e**).

### Reduction of LckCre^+^ GSK-3αβ^fl/fl^ Pre-DP thymocytes expressing TCRβ chain and escape of checkpoints for productive TCRβ-chain rearrangement during β-selection

The proportion of DN thymocytes in 3 wk old LckCre^+^ GSK-3αβ^fl/fl^ mice was significantly higher when compared with LckCre^−^ GSK-3αβ^fl/fl^ littermates suggesting a partial block or delay in transition from the DN to DP stages (13.2% vs. 3.5%) (**Fig. 1a**). Further characterization of DN thymocytes into DN1-DN4 subpopulations was established according to CD117 vs. CD25 expression within the lin^neg^ population as described in Methods. LckCre^+^ GSK-3αβ^fl/fl^ mice were found to have an accumulation of DN3 thymocytes indicating impairment of DN3 to DN4 transition (49% vs. 29.1%). TCRβ chain production did not appear to be compromised at the DN3 stage although there was a small increase in pre-Tα expression (DN3: 10.1% vs. 4.9%) (**Fig. 2a, 2c**).

**Figure 2.**
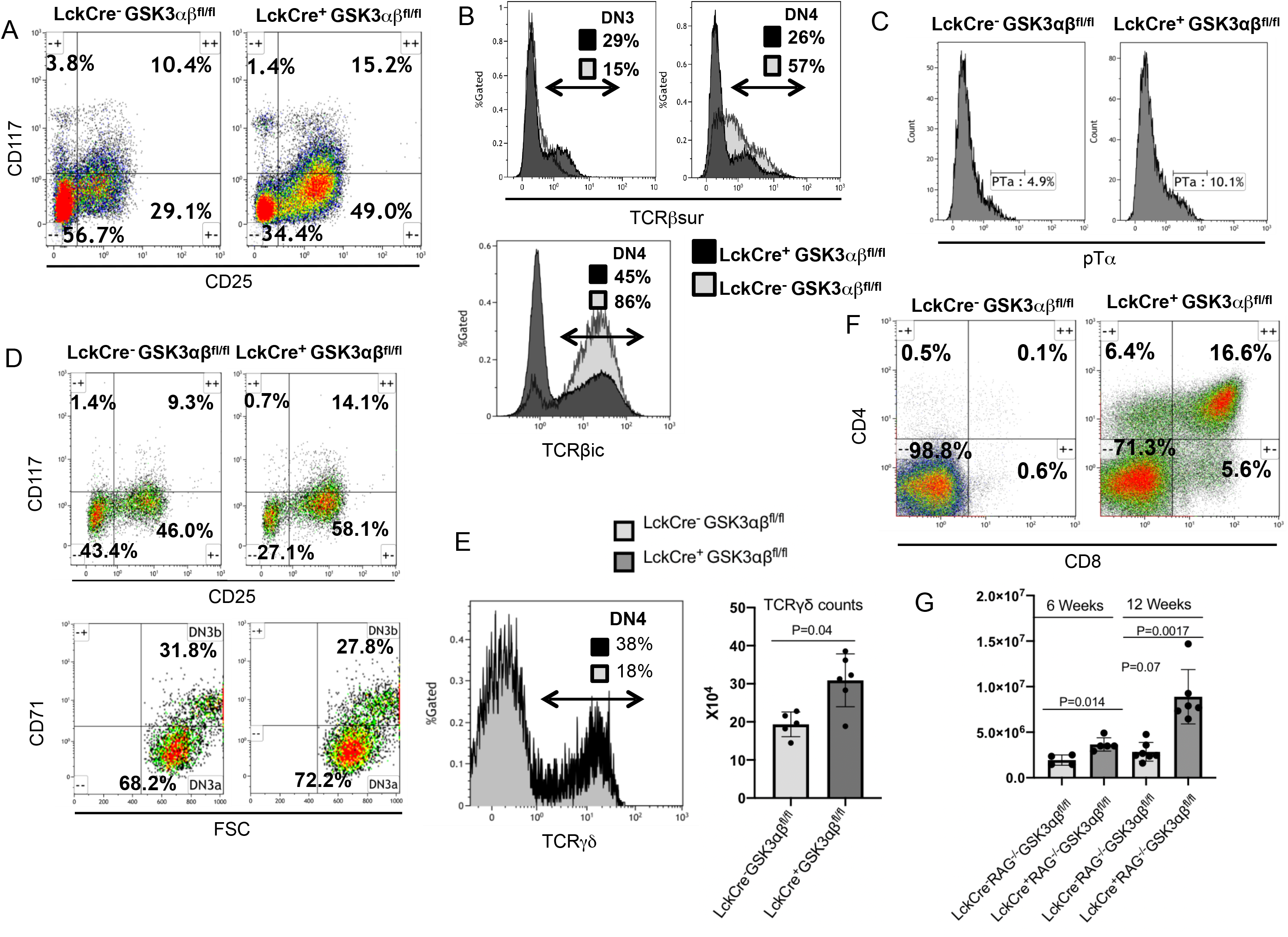
Loss of GSK-3 results in reduced pre TCR surface expression and impaired β-selection. (a) N= 10) LckCre^+^ GSK-3αβ^fl/fl^ and (N=7) LckCre^−^ GSK-3αβ^fl/fl^ mouse thymocytes stained for CD117 vs. CD25 within the DN subset defined as lin^neg^ shows an accumulation of CD117^−^/CD25^+^ (DN3) thymocytes in LckCre^+^ GSK-3αβ^fl/fl^ mice compared to LckCre^−^ GSK-3αβ^fl/fl^ mice and a decrease in DN4 thymocytes (DN3 mean 49.5% +/− 7 vs. 42.4% +/− 5 P= 0.0179 and DN4 35.1% +/− 8.5 vs. 46.3% +/− 2.7 P=.0034). Numbers represent percent cells within quadrants. TCRβ^sur^ expression within DN3 (CD117^−^/CD25^+^/Lin^neg^) sub-population (N=4 mean 26.5% +/− 13.3 vs. 35.9% +/14.1 P= 0.17) and DN4 CD117^−^/CD25^−^/Lin^neg^) sub-population (N=4 mean 26.8% +/− 8 vs. 55.9% +/− 2.8 P= 0.0143). (b) (N=3) Lck^−^ GSK-3αβ^fl/fl^ and (N=4) LckCre^+^ GSK-3αβ^fl/fl^ thymocyte DN4 thymocytes stained for intracellular TCRβ (TCRβic) (mean 81.7% +/− 6.6 vs. 48.6% +/− 13.8 p=0.029). (c) (N=3) LckCre^−^ GSK-3αβ^fl/fl^ and (N=3) LckCre^+^ GSK-3 αβ^fl/fl^ thymocyte sub-population DN3 stained for preTα surface expression. Number represents percent cells positive for preTα (mean 4% +/− 1 vs. 9.17 +/− 0.8 P=0.05 (d) (N= 5) LckCre^−^ GSK-3αβ^fl/fl^ and (N= 6) LckCre^+^ GSK-3αβ^fl/fl^ thymocytes stained for CD71 and gated on DN3 and DN4 sub-populations. CD71 vs. FSC (forward scatter) delineates DN3a from DN3b (DN3a mean 69.7 +/− 3.5 vs. 70.6 +/− 5.1 P=0.42 and DN3b 30.3 +/− 3.5 vs. 29.4 +/− 5.1 P=0.43). (e) (N= 5) LckCre^−^ GSK-3αβ^fl/fl^ and (N=6) LckCre^+^ GSK-3αβ^fl/fl^ DN4 thymocytes stained for TCRγδ (mean 19.1% +/− 3.2% vs. 31% +/6.9% p= 0.015) TCRgd enumeration comparison (P=0.041). (f) (N=6) LckCre^−^ RAG^−/−^ GSK-3αβ^fl/fl^ and (N=5) LckCre^+^ RAG^−/−^ GSK-3αβ^fl/fl^ DP thymocytes from 6 week old mice stained for CD4 and CD8 (mean 0.16% +/− 0.06 vs. 13% +/− 3.3 p=0.004). (g) DP thymocyte numbers for LckCre^−^ RAG^−/−^ GSK-3 αβ^fl/fl^ and LckCre^+^ RAG^−/−^ GSK-3 αβ^fl/fl^ mice at 6 (N=4 P=0.014) and 12 (N=6-7 P=0.017) weeks of age. The difference between 6 and 12 weeks for LckCre^+^ RAG^−/−^ GSK-3 αβ^fl/fl^ mice is P= 0.007. Error bars given as standard error.

Subdivision of DN3 thymocytes into pre-(DN3a) and post-(DN3b) β-selected cells according to CD71 surface expression and forward scatter also failed to reveal any notable impairments during early β-selection as judged by the proportion of DN3b vs. DN3a thymocytes (72.2% : 27.8% vs. 68.2% : 31.8%) (**Fig. 2d**). However, in contrast to DN3 thymocytes, the proportion of LckCre^+^ GSK-3αβ^fl/fl^ DN4 (pre-DP) thymocytes producing and expressing TCRβ chain was greatly reduced (TCRβ^sur^: 26% vs. 57% and TCRβ^ic^: 45% vs. 86%) (**Fig. 2a, 2b**). We note reports showing the DN4 population to harbour NKT cells expressing the invariant Va14Ja18 antigen receptor and it is possible that some residual TCRβ chain expression may be attributed to this population^67^.

Only αβ thymocyte development appeared impaired since expression of γδ TCR was undiminished. In fact, we found the proportion of TCRγδ thymocytes within the DN4 sub-population to be higher for LckCre^+^ GSK-3αβ^fl/fl^ mice than for LckCre^−^ GSK-3αβ^fl/fl^ mice (38% vs. 18%) which is reflected in their increased cell number. This may reflect impaired αβ thymocyte production skewing the normal ratio of αβ to γδ thymocytes, rather than enhancement of γδ T cell production *per se* (**Fig. 2e**).

Given that LckCre^+^ GSK-3αβ^fl/fl^ pre-DP thymocytes had low levels of TCRβ chain production with accumulation of aberrant CD3^+^CD24^−^ CD4^+^CD8^+^ (DP’s), we hypothesized that loss of GSK-3 would allow thymocytes to bypass the strict requirements for productive TCRβ chain rearrangements during β-selection and allow differentiation to DP thymocytes (**Fig. 1b, 2a**). To test this LckCre^+^ RAG1^−/−^ GSK-3αβ^fl/fl^ mice were generated by inter-crossing the LckCre^+^ GSK-3αβ^fl/fl^ mice with RAG1^−/−^ mice. RAG1 (recombination activation gene 1) is required for V(D)J recombination at the TCRβ chain locus^68^. Without RAG1^−/−^ thymocytes cannot assemble the required pre-TCR complex and experience an acute block at the DN3 stage^9^. Here, we show that loss of GSK-3 in the absence of RAG1 was permissive for differentiation from DN3’s to DP thymocytes (16.6% vs. 0.1%) (**Fig. 2f**). We also found an age-dependent increase in thymocyte numbers for LckCre^+^ GSK-3αβ^fl/fl^ RAG1^−/−^ mice (**Fig. 2g**). This finding underscores the extent to which loss of GSK-3 can impact the regulation of β-selection by overriding requirements for competent TCRβ chain rearrangement and pre-TCR dependent differentiation. Of note, a significant proportion of CD4 and CD8 SP’s were also generated which were CD3^+^/CD24^−^, markers normally associated with maturity. However, these cells lacked TCRβ chain surface expression and therefore remained defective (**Fig. 2f and data not shown**).

### LckCre^+^ GSK-3αβ^fl/fl^ DP thymocytes have low levels of productively rearranged TCRβ chain, have high levels of Mcl-1 and are resistant to apoptosis

Levels of LckCre^+^ GSK-3αβ^fl/fl^ DP TCRβ chain rearrangements were reduced 2-3 fold compared to LckCre^−^ GSK-3αβ^fl/fl^ control thymocytes as determined by RT-PCR measurement of Dβ2-Jβ2, Vβ5.1-Jβ2 and Vβ8.2-Jβ2 (**Fig. 3a**). We also assessed the TCR status of DP’s in LckCre^+^ GSK-3αβ^fl/fl^ mice. Thymocytes from 3 wk old mice were stained for surface expression of TCRβ along with CD4 and CD8. Loss of TCRβ expression persisted throughout DP developmental stages, which we defined as CD4^hi^CD8^hi^ (25% vs 66% and median 0.32 vs. 1.43), CD4^lo^CD8^lo^ (45% vs. 95% and median 0.51 vs. 5.58) and CD4^hi^CD8^lo^ (34% vs. 90% and median 0.44 vs. 11.42) respectively (**Fig. 3b**). Together, these data support the view that pre-DP cells in LckCre^+^ GSK-3αβ^fl/fl^ mice bypass normal β-selection checkpoints for proper TCRβ chain rearrangement and expression prior to becoming DP thymocytes.

**Figure 3.**
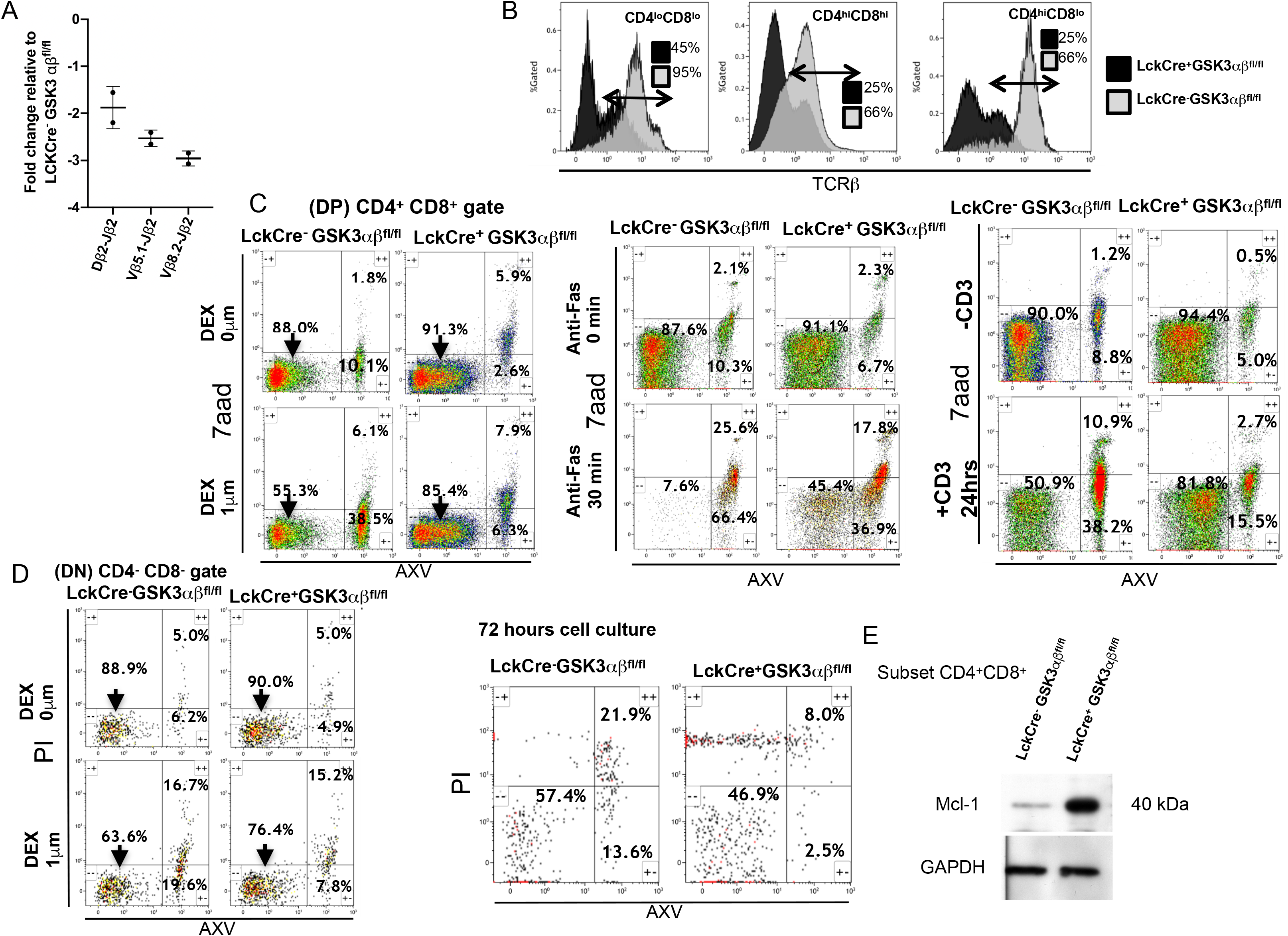
DP thymocytes from Lck^+^ GSK-3 αβ^fl/fl^ mice have compromised TCRβ chain rearrangement, reduced TCR expression and are resistant to cell death. (a) Sorted DP thymocytes from Lck^+^ GSK-3αβ^fl/fl^ and LckCre^−^ GSK-3αβ^fl/fl^ mice subjected to real Time PCR analysis for Dβ2-Jβ2, Vβ5.1-Jβ2 and Vβ8.2-Jβ2 fragment generation as an indication of TCRβ chain rearrangement efficiency. Histograms represent TCRβ chain rearrangements for Lck^+^ GSK-3αβ^fl/fl^ DP thymocytes relative to LckCre^−^ GSK-3αβ^fl/fl^ DP thymocytes. Error bars are given as standard error. (b) Thymocytes from 3 week old N= 6 Lck^−^ GSK-3αβ^fl/fl^ and N= 4 LckCre^+^ GSK-3αβ^fl/fl^ mice were stained for CD4, CD8 and surface TCRβ. DP subsets were analysed for surface TCRβ expression (CD4^lo^/CD8^lo^ mean 82% +/− 9.5 vs. 36.3% +/− 6.1 P= 0.007) and (CD4^hi^/CD8^hi^ mean 53.1 +/− 7 vs. 25.1% +/− 3.8 P= 0.0001) and (CD4^hi^/CD8^lo^ mean 82.9% +/− 13.5 vs. 35.7% +/− 5.3 P= 0.0048). (c,d) Thymocytes from Lck^−^ GSK-3αβ^fl/fl^ and LckCre^+^ GSK-3αβ^fl/fl^ mice were treated as indicated. Cells gated on DP’s and DN’s stained with the cell viability dye 7aad and Anexin V indicates level of apoptosis. Early apoptotic cells are considered 7aad^−^ /AnexinV^+^ while late apoptotic or necrotic cells are 7aad^+^/AnexinV^+^. (e) Sorted DP thymocytes probed with antibody recognizing Mcl-1 with GAPDH as a loading control.

LckCre^+^ GSK-3αβ^fl/fl^ DP thymocytes also exhibited significant resistance to various treatments that induce cell death such as treatment with dexamethasone (1μM for 3 hrs), anti-Fas (30 min), anti-CD3 (24 hrs) and cell culture over a 72 hr period. Early apoptotic and late apoptotic/necrotic cells were identified as Annexin V^+^/7AAD^−^ and Annexin V^+^/7AAD^+^ respectively (dexamethasone induced early apoptosis was 6.3% vs. 38.5%, anti-Fas early apoptosis 36.9% vs. 66.4%, anti-CD3 early apoptosis 15.5% vs. 38.2%). Comparison across these conditions showed that LckCre^+^ GSK-3αβ^fl/fl^ DP thymocytes were significantly resistant to cell death. DN thymocytes of LckCre^+^ GSK-3αβ^fl/fl^ mice were also resistant to apoptotic conditions following dexamethasone treatment (1μM for 3 hrs) (7.8% vs. 19.6%) and cell culture over a 72 hr period (2.5% vs. 13.6%) (**Fig. 3a, 3d and Suppl. Fig. 3b**). Given that GSK-3 is known to regulate stability of the Bcl-2 family survival protein Mcl-1^31–34^ we performed immunoblot analysis on sorted DP thymocytes for Mcl-1 protein levels. LckCre^+^ GSK-3αβ^fl/fl^ DP thymocytes exhibited significantly higher levels of Mcl-1, consistent with enhanced DP cell survival (**Fig. 3e**).

### Rescue of defective positive selection through enforced transgenic TCR expression

Loss of TCR surface expression on thymocyte positive selection intermediates (CD4^hi^CD8^lo^) of the LckCre^+^GSK-3 double knockout mice suggests impaired TCR-mediated signalling during the positive selection process itself (**Fig. 4a**). Normally, CD5 and CD69 expression is directly correlated with signalling through the mature TCR^69^. We observed a significant reduction in CD69 and CD5 expression in the positive selection intermediates (CD4^hi^CD8^lo^) in six wk old LckCre^+^ GSK-3αβ^fl/fl^ mice, a finding consistent with impaired TCR-dependent signalling associated with positive selection^70–73^ (CD69 expression - 21% vs. 83% with median 0.34 vs. 3.19 and CD5 - 16% vs. 88% with median 1.08 vs. 17.73) (**Fig. 4a**). Lack of TCR-mediated signalling may help to explain accumulation of CD4^hi^CD8^lo^ positive selection intermediates as these cells failed positive selection toward mature CD4 and CD8 SP cells (**Fig. 1e**). We also noted a small population of CD24^−^/CD4^+^ and CD24^−^/CD8^+^ SP thymocytes that appear to have relatively high levels of TCRβ and CD3 expression (**data not shown**). This result suggests that it could be the loss of mature TCR expression that contributes to the inability of most LckCre^+^ GSK-3αβ^fl/fl^ DP thymocytes to pass through positive selection to become mature SP cells.

**Figure 4.**
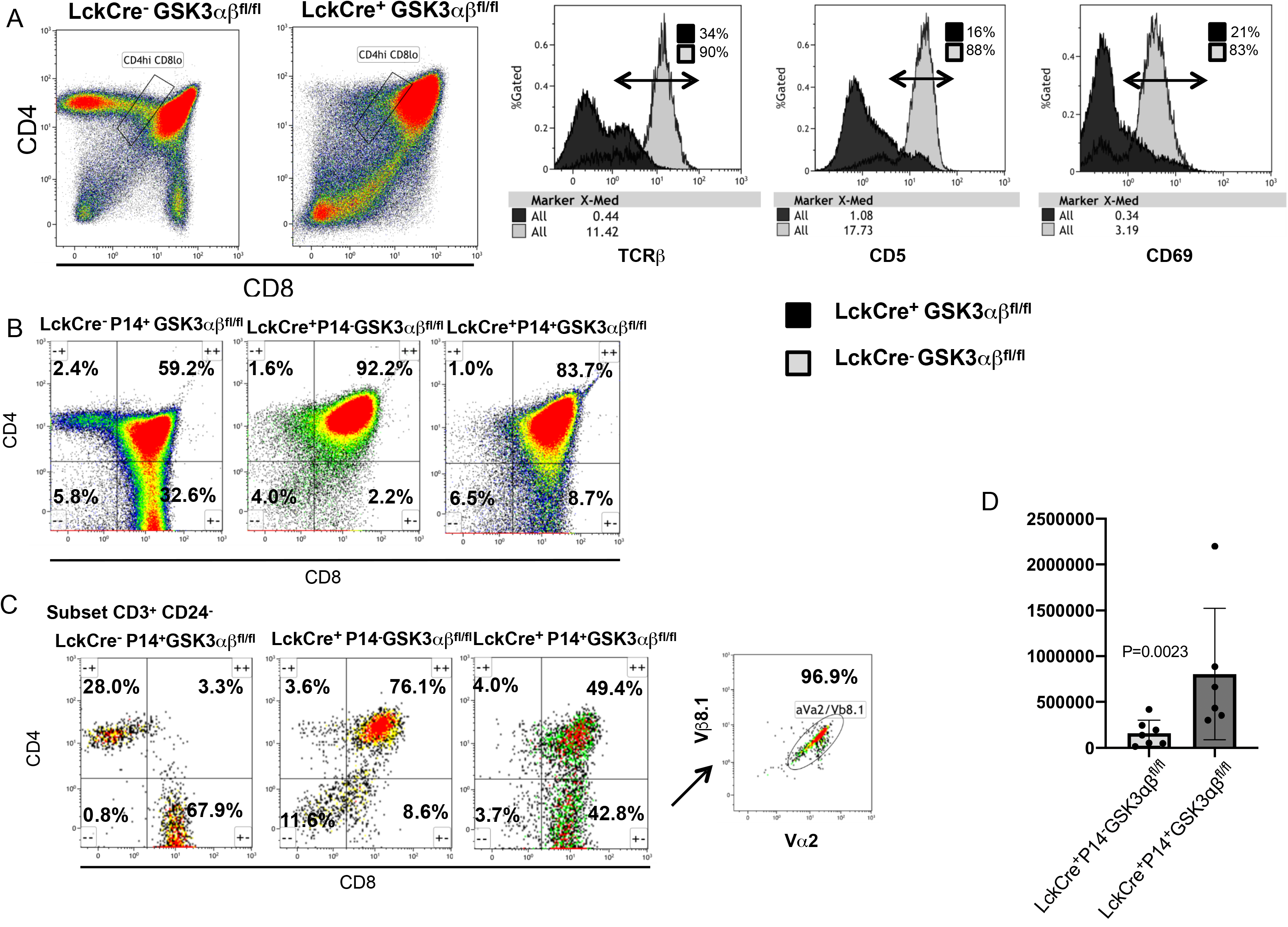
Incorporation of a P14 transgenic TCR alleviates DP to SP selection block (a) Gates indicate CD4^hi^CD8^lo^ positive selection intermediates (CD4^hi^CD8^lo^) from 6 week old Lck^+^ GSK-3αβ^fl/fl^ and LckCre^−^ GSK-αβ^fl/fl^ mice. Reduced levels of surface TCRβ (34% vs. 90% - median 0.42 vs. 11.42), CD69 (21% vs. 83% - median 0.34 vs. 3.19) and CD5 (16% vs. 88% - median 1.08 vs. 17.73) expression for LckCre^+^ GSK-3αβ^fl/fl^ mice was noted indicating reduced TCR signalling during DP to SP selection. (b) Presence of the P14TCR transgene drives differentiation to CD8 SP’s (N=4) LckCre^−^ P14^+^ GSK-3αβ^fl/fl^ (mean 20.63% +/− 10.5), (N=10) LckCre^+^ P14^−^ GSK-3αβ^fl/fl^ (mean 1.2% +/− 1.0) and (N=4) LckCre^+^ P14^+^ GSK-3αβ^fl/fl^ (mean 7.1% +/− 1.2). The difference between LckCre^+^ P14^−^ GSK-3αβ^fl/fl^ and LckCre^+^ P14^+^ GSK-3αβ^fl/fl^ is significant P=0.0029. Proportion of CD8 SP’s within the mature CD3^+^CD24^−^ sub-population (N=4) LckCre^−^ P14^+^ GSK-3αβ^fl/fl^ (mean 43.85% +/− 17, (N=10) LckCre^+^ P14^−^ GSK-3αβ^fl/fl^ (mean 5.2% +/− 2.9) and (N=4) LckCre^+^ P14^+^ GSK-3αβ^fl/fl^ (mean 31.35% +/− 8.2). The difference between LckCre^+^ P14^−^ GSK-3αβ^fl/fl^ and LckCre^+^ P14^+^ GSK-3αβ^fl/fl^ is significant P= 0.001. P14 TCR specificity of CD8’s (CD3^+^ CD24^−^) in LckCre^+^ P14^+^ GSK-3αβ^fl/fl^ mice is confirmed through TCRVβ8.1 and TCRVα2 co-expression (96.9%). (d) Histogram showing cell numbers of mature CD8’s (CD8^+^/CD3^+^/CD24^−^) for littermates LckCre^+^ P14^−^ GSK-3αβ^fl/fl^ and LckCre^+^ P14^+^ GSK-3αβ^fl/f^ mice. The difference is significant P=0.0023. Error bars are indicated as standard error.

To test this idea we used a P14 transgenic TCR model that enforces P14-specific TCR expression and induces positive selection to the CD8 lineage^63^. When supplied with a functional, mature TCR in the form of the P14 transgene, LckCre^+^ P14^+^ GSK-3αβ^fl/fl^ mice produced a greater proportion of CD8 SP’s compared to LckCre^+^ P14^−^ GSK-3αβ^fl/fl^ mice (8.7% vs. 2.2%) (**Fig. 4b**). Further characterization of mature thymocytes (CD3^+^/CD24^−^) in these animals revealed that substantial numbers of mature CD8 SP cells were being generated with expression of the P14 TG (42.8% vs. 8.6%). Co-expression of the Vβ 8.1 and Vα2 T-cell receptors expressed by the P14 transgene confirmed that these CD8 cells were of P14 origin (96.9%). An increase in absolute numbers of CD8^+^/CD3^+^/CD24^−^ cells was also noted for LckCre^+^ P14^+^ GSK-3αβ^fl/fl^ thymocytes compared to LckCre^+^ P14^−^ GSK-3αβ^fl/fl^ thymocytes that lacked the P14 transgene (**Figs. 4b, 4d and suppl. Fig. 3c**). Together these results indicate that the presence of a functional TCR can at least partially rescue the inability of thymocytes lacking GSK-3 to undergo positive selection.

### Thymocytes lacking GSK-3 express high levels of β-catenin with reduced EGR3 and the Notch targets DTX1 and HES1

Given the established role of GSK-3 in suppressing Wnt signalling DP thymocytes were probed for cytoplasmic β-catenin (non-cadherin-associated fraction). High levels of β-catenin were observed only when both GSK-3 isoforms were inactivated. Retention of a single allele of either GSK-3α or GSK-3β resulted in no significant accumulation of β-catenin (**suppl. Fig. 2b and data not shown**). Intracellular staining for β-catenin in the DN3 (lin^neg^CD117^−^CD25^+^), DN4 (lin^neg^CD117^−^CD25^−^), DP (CD4^+^CD8^+^), CD4 (CD4^+^CD8^−^) and CD8 (CD4^−^CD8^+^) sub-populations showed that β-catenin accumulation in the double knocked out thymocytes occurs as early as the DN3 stage and remains high throughout subsequent stages of thymocyte development (**Fig. 5a and suppl. Fig 4a**).

**Figure 5.**
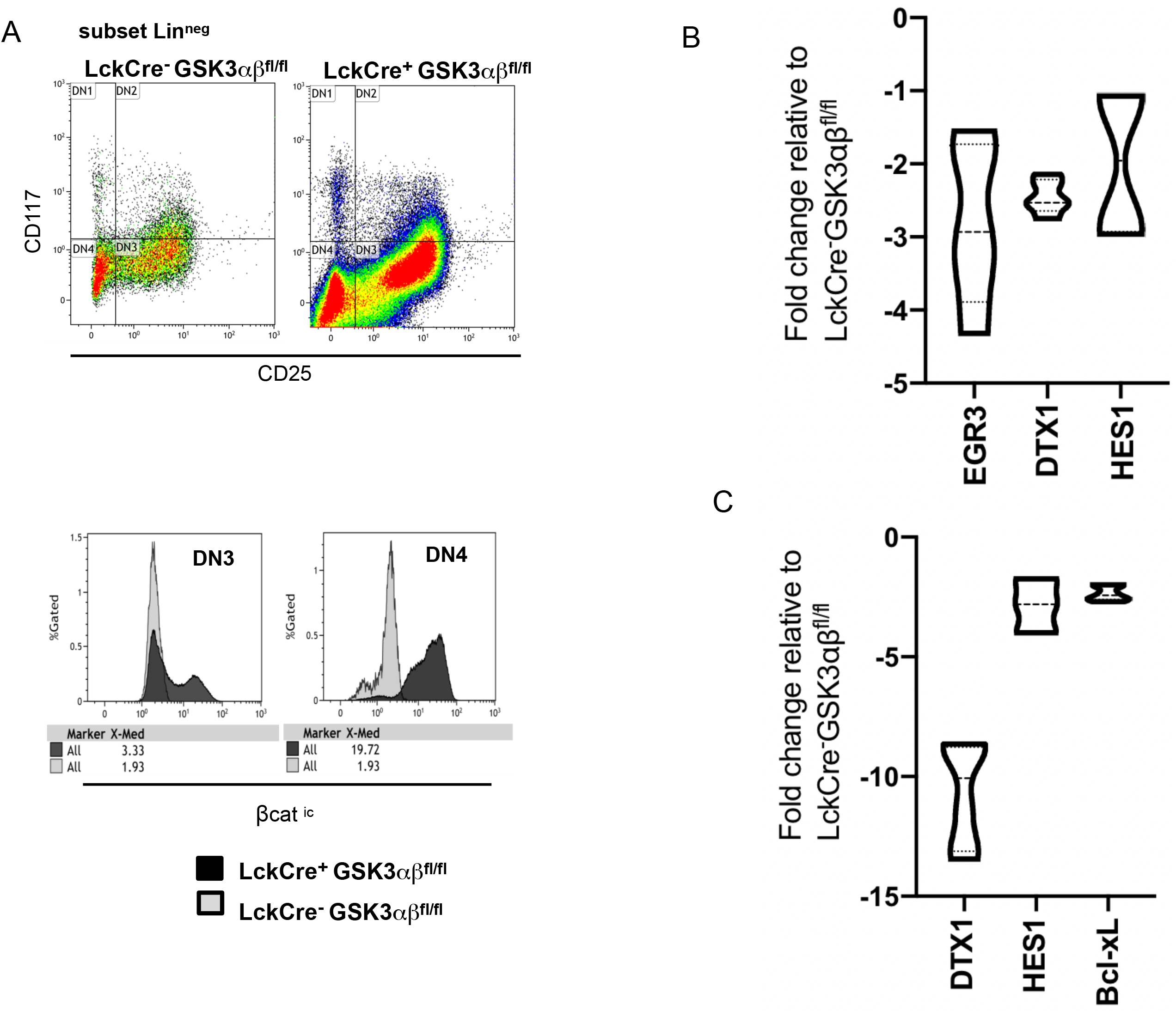
Thymocytes lacking GSK-3 possess high levels of β-catenin with reduced expression of EGR3 and the Notch targets DTX1 and HES1. (a) Intracellular β-catenin expression in DN3 (CD117^−^/CD25^+^/lin^neg^)(median 3.33 vs. 1.93) and DN4 (CD117^−^ /CD25^−^/lin^neg^) (median 19.72 vs. 1.93) sub-populations. (b) Real Time SYBR Green PCR for EGR3, DTX1 and HES1 of sorted LckCre^+^ GSK-3αβ^fl/fl^ and LckCre^−^ GSK-3αβ^fl/fl^ DN4 thymocytes (N=4 experiments with 3 repetitions/reaction/experiment). (c) Real Time SYBR Green PCR for DTX1, HES1 and Bcl-x_L_ and of sorted LckCre^+^ GSK-3αβ^fl/fl^ and LckCre^−^ GSK-3αβ^fl/fl^ DP thymocytes (N=4 experiments with 3 repetitions/reaction/experiment). Transcript levels are plotted as LckCre^+^ GSK-3αβ^fl/fl^ over LckCre^−^ GSK-3αβ^fl/fl^.

β-catenin stabilization has effects on Notch signalling as well as EGR expression^57, 59^. Since both Notch and EGR activity are associated with the pre-DP (DN4) to DP expansion phase following β-selection we used real-time PCR analysis to assess their gene targets in sorted DN4 subpopulations^2, 21, 22, 30^ and observed a 2.5-fold reduction of DTX1, a 2-fold reduction in HES1 and a 3-fold reduction of Egr3 transcripts in LckCre^+^ GSK-3αβ^fl/fl^ DN4 thymocytes compared to control littermates. These findings are consistent with the loss of pre-DP proliferation and DP hypocellularity seen in 3 wk old LckCre^+^ GSK-3αβ^fl/fl^ mice (**Fig. 1d and Fig. 5b**). Notch activity, as indicated by DTX1 and HES1 transcript levels, was also significantly reduced in LckCre^+^ GSK-3αβ^fl/fl^ DP thymocytes (10- and 2-fold, respectively). Since we observed enhanced DP survival and an increase in the Bcl-2 family survival protein Mcl-1, we also examined Bcl-X_L_, another Bcl-2 family member that has been linked to DP survival, but found no significant change (**Fig. 5c and data not shown**).

### LckCre^+^ GSK-3αβ^fl/fl^ mice produce anergic peripheral T cells and succumb to lymphomas

Peripheral lymphoid splenic CD4 and CD8 SP’s were significantly reduced in both percentage of population and cell number in GSK-3 knock-out mice but expressed high levels of CD3 and TCR (**Fig. 6a, 6b and suppl. Fig. 4b**). However, these lymphocytes failed to proliferate in response to either anti-CD3 +/− IL-2 or anti CD3/CD28 +/− IL-2 mediated stimulation, as measured by the CFSE dye incorporation, indicating TCR mediated signaling is impeded (**Fig. 6c and data not shown**). LckCre^+^ GSK-3αβ^fl/fl^ mice developed peripheral lymphomas with a penetrance of 77% and reached endpoint at a median of 157 days in comparison to LckCre^−^GSK-3αβ^fl/fl^ and LckCre^+^ control mice, which showed no significant decrease in survival within a comparable time period. LckCre^+^RAG^−/−^GSK-3αβ^fl/fl^ mice had a median endpoint survival of 131 days with a penetrance of 43% and therefore do not show the enhanced survival observed for stabilized β-catenin mice within a RAG^null^ background^60^ (**Fig. 6d**). Furthermore, unlike stabilized β-catenin, thymic lymphomas occurred with much lower frequency, presented heterogeneous CD4 and CD8 expression and exhibited high levels of Mcl-1 (**Fig. 6e and data not shown**). Peripheral lymphomas also had heterogeneous CD4 and CD8 expression and were determined to be oligoclonal through PCR analysis of the 22 TCRVβ clone families within the CDR3 region, in addition to screening for overrepresentation of TCRVβ gene families by flow cytometry (**Fig. 6f, suppl. Fig. 4c and data not shown**).

**Figure 6.**
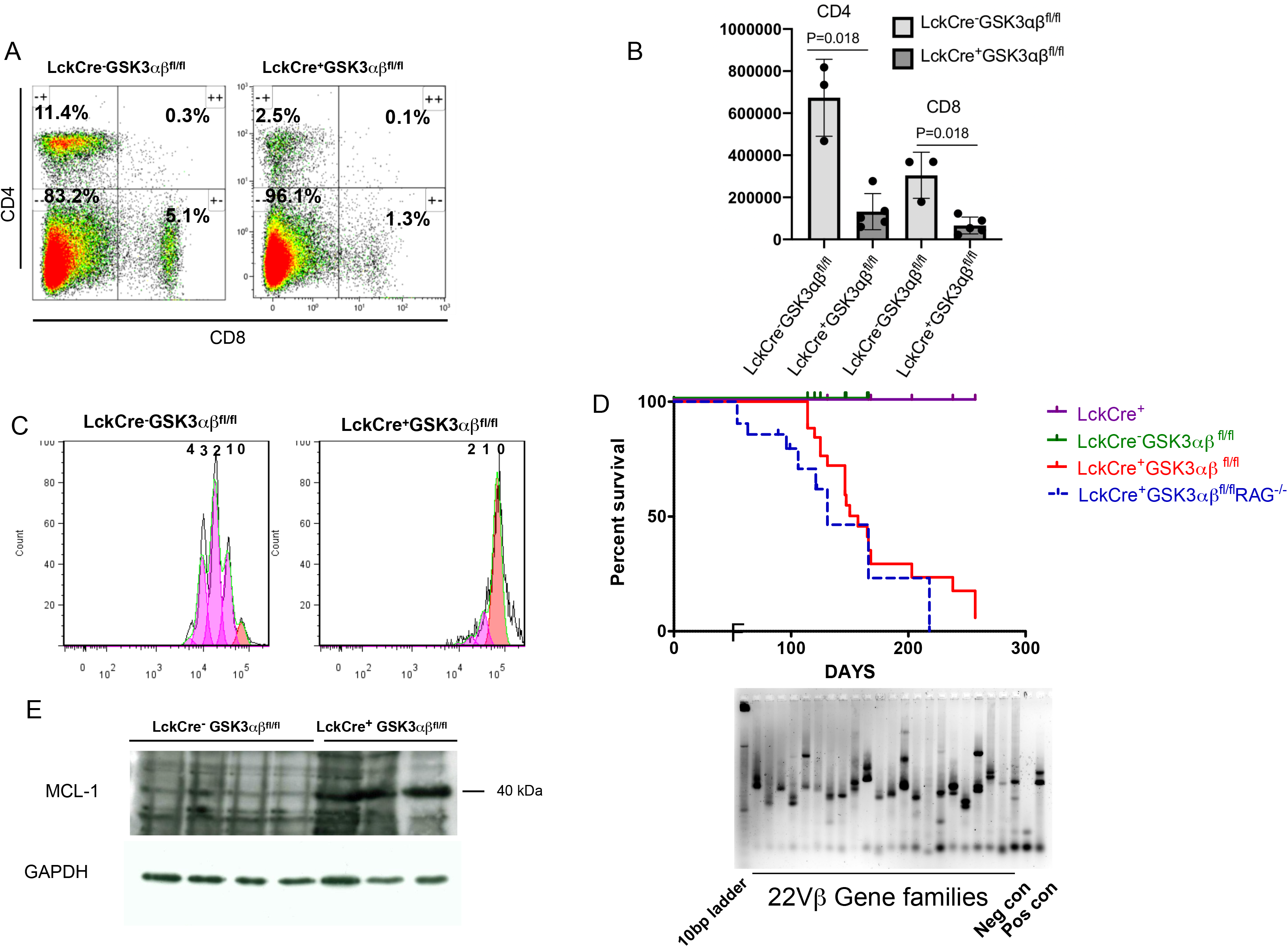
LckCre^+^ GSK-3αβ^fl/fl^ mice peripheral T cells unresponsive to stimulation and succumb to lymphomas with high levels of Mcl-1 expression. (a) Peripheral CD4 and CD8 SP Splenocytes from (N=3) LckCre^−^GSK-3αβ^fl/fl^ and (N=5) LckCre^+^GSK-3αβ^fl/fl^ mice stained with CD4 and CD8 antibodies (CD4 mean 11.5% +/− 2.0 vs. 2.3% +/− 0.8 P=0.0179, CD8 mean 5.2% +/− 1.6 vs. 1.2% +/− 0.25 P=0.0179. (b) CD4 and CD8 cell numbers were calculated and one tailed non-paired t-tests performed. LckCre^+^GSK-3αβ^fl/fl^ mice have significantly less CD4 (P<0.0179) and CD8 (P<0.0179) splenic cell numbers. (c) Proliferative response splenocytes from LckCre^−^GSK-3αβ^fl/fl^, and LckCre^+^GSK-3αβ^fl/fl^ mice were loaded with CFSE for population tracking and stimulated with anti-CD3 and IL-2 for 72 hours. Proliferation modeling was carried out on CD4 SP’s using the proliferation platform of FlowJo software (Treestar). This provided comparative information on the division index, proliferation index and the percent divided. LckCre^−^GSK-3αβ^fl/fl^ (division index = 1.33, proliferation index = 1.65, divided =80.56%) and LckCre^+^GSK-3αβ^fl/fl^ (division index = 0.13, proliferation index = 1.18, divided = 10.66%). Similar results were obtained using anti-CD3 + anti-CD28 +/− IL-2 (data not shown). (d) Kaplan-Meier survival curves of (N= 17) LckCre^+^, (N=15) LckCre^−^ GSK-3αβ^fl/fl^, (N=18) LckCre^+^GSK-3αβ^fl/fl^ and (N=21) LckCre^+^RAG^−/−^GSK-3αβ^fl/fl^ mice. LckCre^+^GSK-3αβ^fl/fl^ and LckCre^+^RAG^−/−^GSK-3αβ^fl/fl^ mice reached endpoint at a median 157 and 131 days respectively with a penetrance of 77% and 43% while LckCre^+^, and LckCre^−^GSK-3αβ^fl/fl^ control mice did not reach endpoint during comparable time periods. Log-rank scores indicated that there were significant differences in the survival curves of LckCre^+^GSK-3αβ^fl/fl^ and LckCre^+^RAG^−/−^GSK-3αβ^fl/fl^ mice (P<0.0001). (e) Mcl-1 immunoblot of thymocytes from 14 wk old (N=4) LckCre^−^GSK-3αβ^fl/fl^ and (N=3) LckCre^+^GSK-3αβ^fl/fl^ animals (presenting thymic lymphoma).(f) TCR Vβ repertoire clone screen of CD8 sorted splenocytes taken from a LckCre^+^GSK-3αβ^fl/fl^ mouse with a peripheral lymphoma. Prominence of single bands instead of smears in TCR Vβ lanes indicates oligoclonal nature of lymphoma.

## Discussion

Given the central role GSK-3 in mediating PI3K and Wnt signals along with implication in a variety of signaling pathways which impact β-selection, the goal of this study was to identify steps in early T cell development and beyond affected by loss of GSK-3^13–20; 25, 75^. Indeed, pharmacological inhibition of GSK-3 within an *in vitro*, stromal-free cell culture system have suggested that DP thymocytes could be produced in the absence of preTCR and Notch1-mediated signalling^16^.

Here, we found that the normally strict requirements for successful β-selection are circumvented by inactivation of GSK-3. During thymocyte ontogeny, β-selection is a crucial checkpoint during which early thymocytes expressing a repertoire of newly rearranged TCRβ chains of varying specificities and avidities are selected following proper TCR β-chain rearrangement. A series of studies have shown signals downstream of the pre-TCR and Notch to converge on β-selection to provide the appropriate survival, proliferation and differentiation signals necessary for generating a large pool of competent DP cells upon which appropriate selection to mature T cell lineages may occur^23^. Perturbations during this early checkpoint can lead to genomic instability and aberrant development, which can lead to lymphomagenesis^7^.

We observed a block during the DN3-DN4 transition in LckCre^+^ GSK-3αβ^fl/fl^ mice with significant loss of TCRβ chain from both the surface and inside the cell, but without loss of TCRgd expression or numbers (**Fig 2a, b and e**). Furthermore, Notch targets DTX and HES1 were reduced suggesting lower Notch1 activity, followed by a four-fold reduction in DP cellularity in 3 wk old LckCre^+^ GSK-3αβ ^fl/fl^ mice (**Fig 5b and Fig 1d**). Resulting DP thymocytes expressed significantly less TCRβ chain along with a reduction in generation of productive TCRβ V(D)J fragments (**Fig 3a and b**). It has been reported that β-selected DN3b and DN4 cells can forgo proliferation and differentiate into DP cells in absence of Notch signals *in vitro*^2, 78^. Taken together, one possibility is that the strict requirements for productive TCRβ chain expression during β-selection are bypassed promoting aberrant differentiation without prerequisite clonal expansion of β- selected precursors. Consistent with this loss of proliferative burst was an observed decrease in EGR3 expression in pre-DP DN4 cells, since EGR3 is known to play a proliferative role during β-selection (**Fig 5b**)^21, 22^. The fact that DN thymocytes show elevated survival argues against the idea that loss of DP cellularity is due to enhanced cell death during transition (**Fig 3d**).

Along with pre-DP self-renewal and expansion, survival signals are necessary before thymocytes can complete transition to DP’s. Several studies have shown components of PI3K signaling to be important for pre-TCR mediated survival, relevant since GSK-3 is a direct target of PI3K-PKB/AKT signaling^24, 25, 29^. Specifically, loss of various combinations of PKB/AKT isoforms has been shown to affect pre-TCR mediated survival and result in DP hypocellularity due to elevated apoptosis, while ablation of PTEN, a negative regulator of PKB/AKT, has the opposite effect. Furthermore, studies employing a knock-in mutation of PDK1 (PDK1^L155E^), which is restricted to signaling through PKB/AKT, suggest that PDK1-mediated differentiation, but not proliferation, signals through PKB/AKT during β-selection^26–29, 79^. Supporting this finding, DN3 cells expressing an activated mutant of PKB/AKT were shown to have elevated survival and to not require Notch signals for progression to the DP stage using the OP-9 cell culture system. Constitutively active Akt1 (*Myr-Akt1*) was also able to rescue DP production in RAG^null^ mice, which have curtailed TCRβ-chain rearrangements and therefore do not possess a functional pre-TCR^26, 27^. To examine whether LckCre^+^GSK-3αβ ^fl/fl^ pre-DP’s could escape requirements for pre-TCR expression during β-selection we tested the effect of a *RAG1^null^* mutation which displays a severe block in pre-DP-DP transition^10, 80^. LckCre^+^ GSK-3αβ^fl/fl^ RAG1^null^ mice exhibited robust differentiation to DP’s despite impaired TCRβ-chain rearrangement suggesting that LckCre^+^ GSK-3αβ^fl/fl^ pre-DP’s can bypass the normal requirements of pre-TCR expression during β-selection (**Fig 2f**). We observed a similar age-dependent increase in LckCre^+^ GSK-3αβ^fl/fl^ RAG1^null^ thymocytes to that found in activated PKB/Akt *RAG1^null^* mice (**Fig 2g**).

Sen and colleagues have shown that β-catenin is transiently expressed during β- selection and that it facilitates pre-TCR signalling. They concluded that under normal circumstances β-catenin expression is tightly regulated and must be down-regulated following β-selection, otherwise sustained expression impairs development through oncogene-induced senescence followed by p53-dependent apoptosis, which, they suggest, contributes to reduced DP cell numbers observed in β-catenin-Tg mice^57, 81^. Successful double-strand break repair is required during TCRβ V(D)J recombination and is dependent upon p53 to mediate cell cycle arrest required for productive V(D)J rearrangement. Loss of p53 function in this context has been associated with genomic instability and lymphomagenesis in both mice and humans, along with DN-DP transition^17, 82^. Interestingly GSK-3 is known to regulate p53 activity through phosphorylation of the Tip60 histone acetyltransferase^18^. It is therefore tempting to speculate that loss of GSK-3 may contribute to escape of LckCre^+^ GSK-3αβ^fl/fl^ preDP’s from β-selection through lack of p53-mediated cell cycle arrest and apoptosis, thereby contributing to lymphomagenesis.

Given the potential of GSK-3 to mediate survival and our observation that DP cellularity in LckCre^+^ GSK-3αβ^fl/fl^ mice rises substantially by seven weeks, we noted substantial resistance of these DP cells to apoptotic stimuli, likely contributing to their increased number over time (**Fig 3c**). This contrasts with other models that exhibit high levels of β-catenin, notably Ctnnb1^loxpex3^, and Apc^lox/lox468^, which each show continued DP hypocellularity attributed to increased apoptosis^59,60^. In keeping with our findings of enhanced LckCre^+^ GSK-3αβ^fl/fl^ DP cell survival, we found high levels of the Bcl-2 family survival protein Mcl-1 (**Fig 3e**). Mcl-1 is normally expressed throughout thymocyte development and conditional deletion of Mcl-1 impairs DN/DP survival. It has also been suggested that MCL-1 contributes to pre-TCR-mediated survival^31, 32^. Mcl-1 stability can be regulated via GSK-3, which supports the idea that survival of LckCre^+^ GSK-3αβ^fl/fl^ DP cells is GSK-3-dependent and can override pro-apoptotic signals derived from β-catenin stabilization^35^.

We noted that although DP cells were generated in LckCre^+^ GSK-3αβ^fl/fl^ mice, few mature CD4 or CD8 SP’s resulted and their numbers did not increase as the DP thymocyte population rose over time (**Fig 1a, b, c, and Suppl Fig 2c**). Loss of mature CD4 or CD8 SP’s was also evident in peripheral lymphoid organs and these cells failed to respond to TCR stimulation, even though they expressed relatively high levels of TCR and CD3 (**Fig 6a, b and Suppl. Fig 4b**). Given that positive selection requires appropriate levels of mature αβTCR it is not surprising that DP thymocytes of LckCre^+^ GSK-3αβ^fl/fl^ mice fail to produce significant SP numbers and accumulate as positive selection intermediates (**Fig 1e**). Previous studies have indicated that deregulation of both β-catenin and Notch can impact DP-SP differentiation and we found significantly less Notch activity, along with high levels of β-catenin, throughout DP development including the positive selection intermediate stage. Furthermore, β-catenin has been shown to be post-transcriptionally regulated by TCR signaling through PI3K involving GSK-3 (**Fig 5c and Suppl. 4a**)^46, 83, 84^. Accumulated TCR^lo^CD4^hi^CD8^lo^ positive selection intermediates lacked CD5 or CD69 expression, which is a further indication of compromised TCR signalling. This suggests that while positive selection may be initiated, TCR functions are impaired resulting in failure of DP’s to undergo positive selection. However, we did find that, while significantly reduced in number, some CD4^+^CD24^−^ and CD8^+^CD24^−^ SP thymocytes were generated in LckCre^+^ GSK-3αβ^fl/fl^ mice with relatively high levels of mature TCR as measured through TCR β chain and CD3 expression (data not shown). We therefore tested the possibility that enforced TCR expression might restore aspects of positive selection blocked by loss of GSK-3 by using a transgenic P14 cassette used in previous studies of positive selection to supply a functional TCR^73^. This TCR cassette directs positive selection of CD8 SP’s and generated a significant proportion of P14-specified mature CD8 SP thymocytes in LckCre^+^ P14^+^ GSK-3αβ^fl/fl^ mice. We observed only modest restoration of SP CD8 numbers in LckCre^+^ P14^+^ GSK-3αβ^fl/fl^ mice but this may be attributed to a documented lack of engineered TCR’s ability to fully restore the proliferative burst associated with pre-DP expansion^7, 25, 85, 86^.

In summary, GSK-3 plays multiple roles in T cell differentiation, several of which are via Wnt/β-catenin but clearly also via other mechanisms including Mcl-1 and mediation of PI3K signalling.

## Acknowledgements

This work was supported a Foundation grant from the Canadian Institutes of Health Research to JRW. We also acknowledge support from the flow cytometry core facilities at the University Health Network and Lunenfeld-Tanenbaum Research Institute, Toronto, Ontario.

## Supplementary figure legends

**Suppl. Fig. 1**. Strategy to create conditional knockout of GSK-3α and GSK-3β genes. Targeting strategy for GSK-3α. After homologous recombination with the targeting vector, exon 2 was replaced with a LoxP-flanked (floxed) exon 2 and FRT-flanked neomycin-resistance cassette. B, BamHI; A, ApaI. Targeting strategy for GSK-3β. After homologous recombination, exon 2 was replaced with a neomycin-resistance cassette and single FRT site. B, BglII. Asterisks represent hybridization sites for Southern blot probes.

**Suppl. Fig 2**. A. Conditional loss of GSK-3α and GSK-3β isoforms in mouse thymocytes. Genomic DNA analysed by PCR for LckCre-mediated excision of GSK-3α and GSK-3β exon 2 in thymocytes of LckCre^+^ GSK-3αβ^fl/fl^ and LckCre-GSK-3 αβ^fl/fl^ mice sorted into DN3, DN4, CD4^+^/CD8^+^, CD4^+^/CD8^−^ and CD4^−^/CD8^+^ sub-populations.B. Immunoblot of LckCre^+^ GSK-3 α^fl/fl^ β^fl/wt^ (α^3/4^), LckCre^+^ GSK-3 α^fl/wt^ β^fl/fl^ (β^3/4^), LckCre^+^ GSK-3αβ^fl/fl^ and LckCre^−^ GSK-3αβ^fl/fl^ sorted DP thymocytes probed with antibodies recognizing both GSK-3α and GSK-3β isoforms, β-catenin and GAPDH as a loading control. C. LckCre^−^ GSK-3αβ^fl/fl^ and LckCre^+^ GSK-3 αβ^fl/fl^ thymocytes of 6 wk old mice stained for CD4 vs. CD8 expression along with a CD4 vs. CD8 stain of mature T cell sub-populations gated on the CD3^+^/CD24^−^ subset.

**Suppl. Fig. 3**. A. LckCre^+^ GSK-3α^fl/fl^ β^fl/wt^ (N=3), LckCre^+^ GSK-3α^fl/wt^ β^fl/fl^ (N=3) and LckCre^−^ GSK-3α^fl/fl^ β^fl/wt^ (N=3) thymocytes of 3 wk old mice were stained for CD4 vs. CD8 expression along with a CD4 vs. CD8 stain of mature T cell subpopulations gated on the CD3^+^/CD24^−^ subset. Comparisons show no significant difference between populations (CD4^+^ P=0.51, CD8^+^ P=0.58, CD4^+^CD3^+^CD24^−^ P= 0.12 and CD8^+^CD3^+^CD24^−^ P=0.13). B. Thymocytes from LckCre^−^ GSK-3αβ^fl/fl^ and LckCre^+^ GSK-3αβ^fl/fl^ mice were placed in culture for the times indicated. Cells were gated on the CD4^+^ CD8^+^ (DP) sub-population and stained with the cell viability dye 7aad and Annexin V (AXV) to indicate level of apoptosis. Early apoptotic cells are considered 7aad-/Annexin V^+^ while late apoptotic or necrotic cells are 7aad^+^/Annexin V^+^. C. Presence of the P14TCR transgene drives differentiation to CD8 SP’s (N=4) LckCre^−^ P14^+^ GSK-3αβ^fl/fl^ (mean 20.63% +/− 10.5), (N=10) LckCre^+^ P14^−^ GSK-3αβ^fl/fl^ (mean 1.2% +/− 1.0) and (N=4) LckCre^+^ P14^+^ GSK-3αβ^fl/fl^ (mean 7.1% +/− 1.2). The difference between LckCre^+^ P14^−^ GSK-3αβ^fl/fl^ and LckCre^+^ P14^+^ GSK-3αβ^fl/fl^ is significant P=0.0029. Proportion of CD8 SP’s within the mature CD3^+^CD24^−^ sub-population (N=4) LckCre^−^ P14^+^ GSK-3αβ^fl/fl^ (mean 43.85% +/− 17, (N=10) LckCre^+^ P14^−^ GSK-3αβ^fl/fl^ (mean 5.2% +/− 2.9) and (N=4) LckCre^+^ P14^+^ GSK-3αβ^fl/fl^ (mean 31.35% +/− 8.2). The difference between LckCre^+^ P14^−^ GSK-3αβ^fl/fl^ and LckCre^+^ P14^+^ GSK-3αβ^fl/fl^ is significant P= 0.001. P14 TCR specificity of CD8’s (CD3^+^ CD24^−^) in LckCre^+^ P14^+^ GSK-3αβ^fl/fl^ mice is confirmed through TCRVβ8.1 and TCRVα2 co-expression (96.9%). TCRVβ8.1 vs. TCRVα2 co-expression shows that majority of mature CD8 SP’s are specified by the P14 transgene in LckCre^+^ P14^+^ GSK-3αβ^fl/fl^ and LckCre^−^ P14^+^ GSK-3αβ^fl/fl^ mice while the P14 transgene cannot be detected in LckCre^+^ P14^−^ GSK-3αβ^fl/fl^ CD3^+^CD24^−^CD8 SP’s.

**Suppl. Fig. 4**. A. Intracellular β-catenin expression expressed as median fluorescent intensity within gated thymocyte sub-populations. B. Splenic CD4^+^ and CD8^+^ SP populations from LckCre^+^ GSK-3αβ^fl/fl^ mice had comparable levels of TCR and CD3 expression to LckCre^−^ GSK-3αβ^fl/fl^ mice even though numbers were significantly reduced. C. FACS of 5 individual LckCre^+^ GSK-3αβ^fl/fl^ lymphomas reveal heterogeneity as judged by CD3, CD4, CD8, and CD19 markers.

