## Supplemental figures for "Regulation of thymocyte β-selection, development and positive selection by glycogen synthase kinase-3"

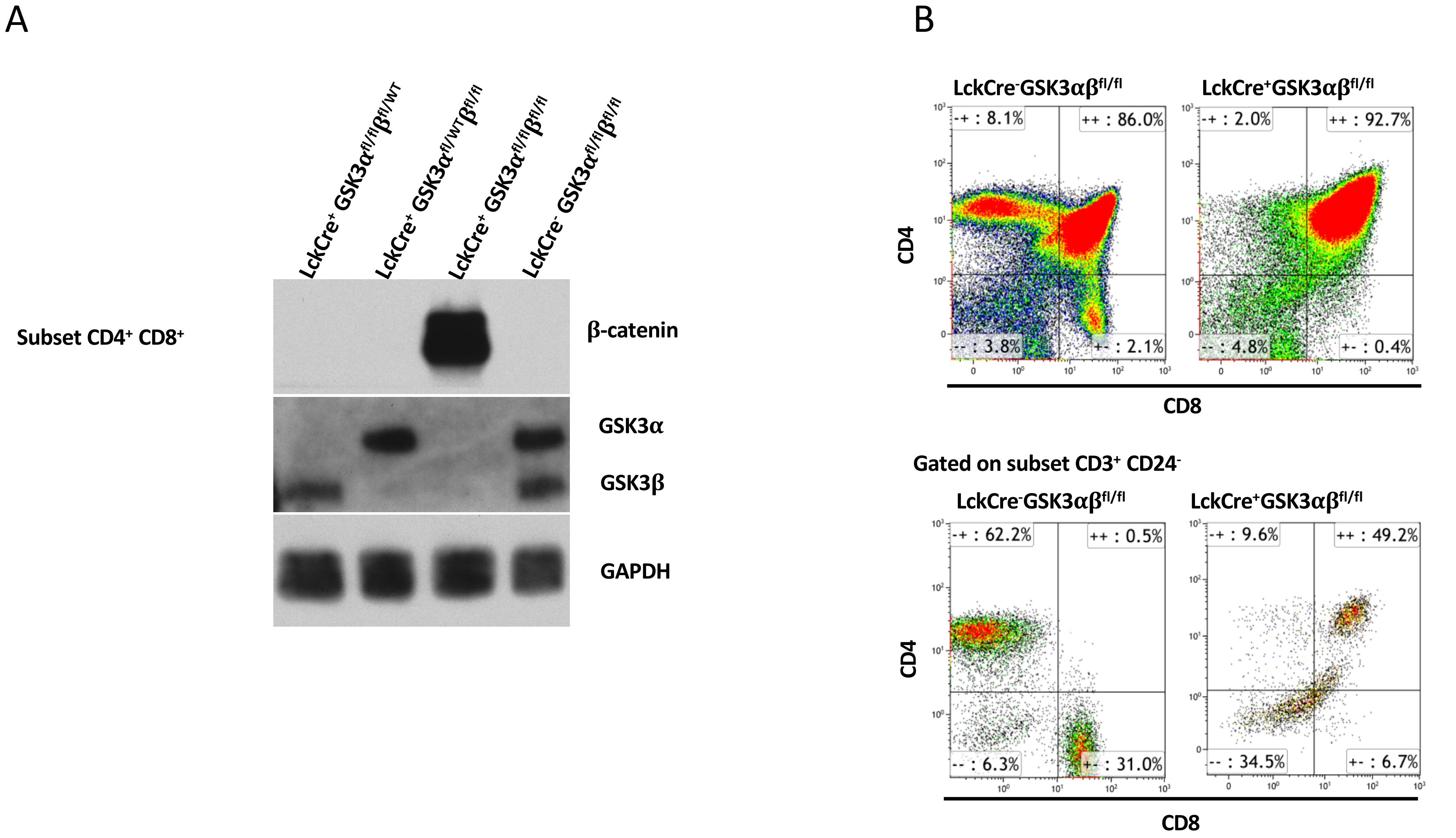


**Suppl. Fig 1.** A. Immunoblot of LckCre$^+$ GSK-3α$^{fl/fl}$ β$^{fl/wt}$ (α$^{3/4}$), LckCre$^+$ GSK-3α$^{fl/wt}$ β$^{fl/fl}$ (β$^{3/4}$), LckCre$^+$ GSK-3αβ$^{fl/fl}$ and LckCre$^-$ GSK-3αβ$^{fl/fl}$ sorted DP thymocytes probed with antibodies recognizing both GSK-3α and GSK-3β isoforms, β-catenin and GAPDH as a loading control. C. LckCre$^-$ GSK-3αβ$^{fl/fl}$ and LckCre$^+$ GSK-3αβ$^{fl/fl}$ thymocytes of 6 week old mice stained for CD4 vs. CD8 expression along with a CD4 vs. CD8 stain of mature T cell sub-populations gated on the CD3$^+$/CD24$^-$ subset.

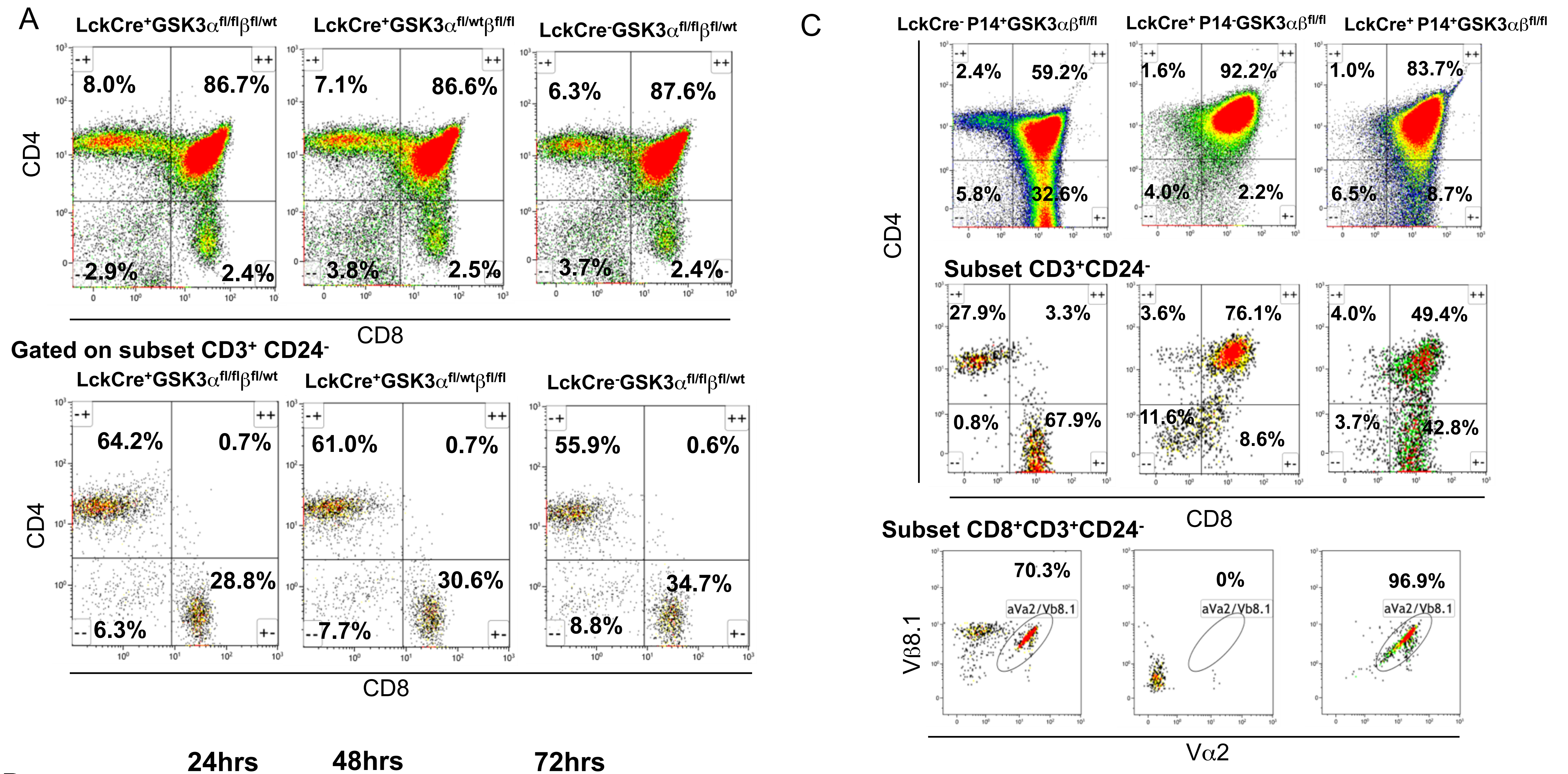

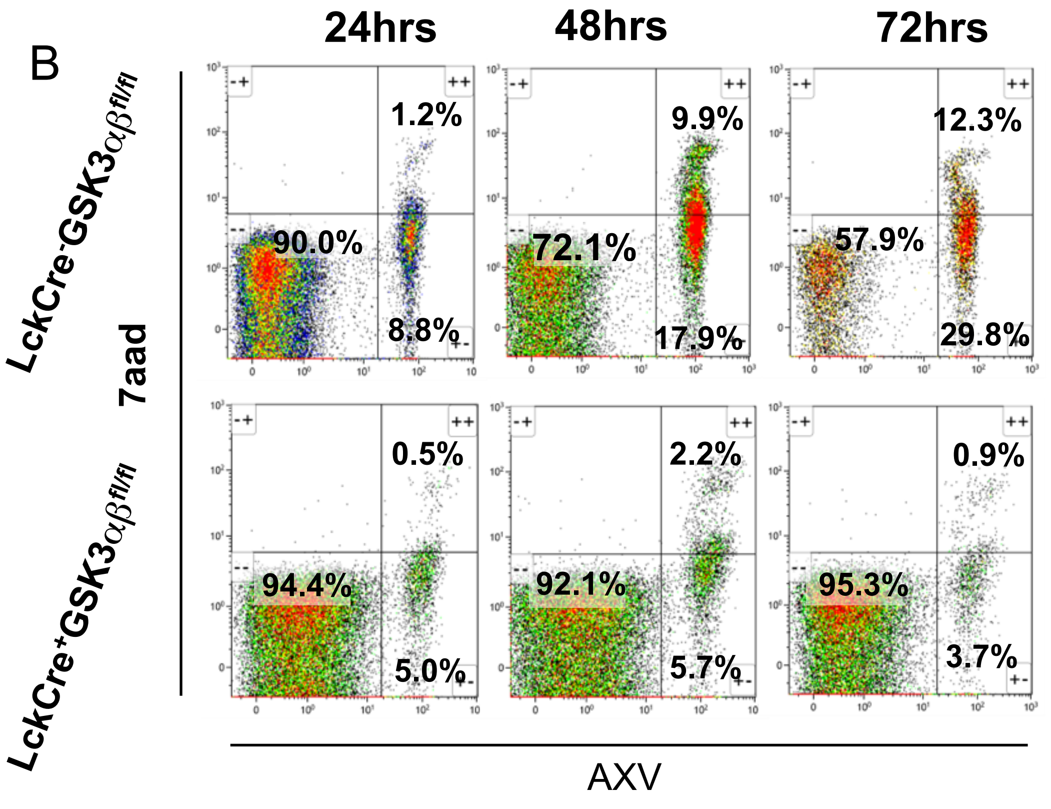


**Suppl. Fig. 2.** A. LckCre$^{+}$ GSK-3α$^{fl/fl}$ β$^{fl/wt}$ (N=3), LckCre$^{+}$ GSK-3α$^{fl/wt}$ β$^{fl/fl}$ (N=3) and LckCre$^{-}$ GSK-3α$^{fl/fl}$ β$^{fl/wt}$ (N=3) thymocytes of 3 wk mice stained for CD4 vs. CD8 expression along with a CD4 vs. CD8 stain of mature T cell subpopulations gated on CD3$^{+}$/CD24$^{-}$ subset. No significant difference between populations (CD4$^{+}$ P=0.51, CD8$^{+}$ P=0.58, CD4$^{+}$CD3$^{+}$CD24$^{-}$ P= 0.12 and CD8$^{+}$CD3$^{+}$CD24$^{-}$ P=0.13). B. Thymocytes from LckCre$^{-}$ GSK-3αβ$^{fl/fl}$ and LckCre$^{+}$ GSK-3αβ$^{fl/fl}$ mice cultured for times indicated. Cells gated on the CD4$^{+}$ CD8$^{+}$ (DP) sub-population and stained with 7aad and Annexin V (AXV) to indicate apoptosis. Early apoptotic cells are considered 7aad-/Annexin V$^{+,}$ late apoptotic or necrotic cells are 7aad$^{+}$/Annexin V$^{+}$. C. P14TCR transgene drives differentiation to CD8 SP's (N=4) LckCre$^{-}$ P14$^{+}$ GSK-3αβ$^{fl/fl}$ (mean 20.63% +/-10.5), (N=10) LckCre$^{+}$ P14$^{-}$ GSK-3αβ$^{fl/fl}$ (mean 1.2% +/- 1.0) and (N=4) LckCre$^{+}$ P14$^{+}$ GSK-3αβ$^{fl/fl}$ (mean 7.1% +/- 1.2). Difference between LckCre$^{+}$ P14$^{-}$ GSK-3αβ$^{fl/fl}$ and LckCre$^{+}$ P14$^{+}$ GSK-3αβ$^{fl/fl}$ is significant P=0.0029. Proportion of CD8 SP's within the mature CD3$^{+}$CD24$^{-}$ sub-population (N=4) LckCre$^{-}$ P14$^{+}$ GSK-3αβ$^{fl/fl}$ (mean 43.85% +/- 17, (N=10) LckCre$^{+}$ P14$^{-}$ GSK-3αβ$^{fl/fl}$ (mean 5.2% +/-2.9) and (N=4) LckCre$^{+}$ P14$^{+}$ GSK-3αβ$^{fl/fl}$ (mean 31.35% +/- 8.2). Difference between LckCre$^{+}$ P14$^{-}$ GSK-3αβ$^{fl/fl}$ and LckCre$^{+}$ P14$^{+}$ GSK-3αβ$^{fl/fl}$ is significant P=.001. P14 TCR specificity of CD8's (CD3$^{+}$ CD24$^{-}$) in LckCre$^{+}$ P14$^{+}$ GSK-3αβ$^{fl/fl}$ mice confirmed via TCRVβ8.1 and TCRVα2 co-expression (96.9%). TCRVβ8.1 vs. TCRVα2 co-expression shows majority of mature CD8 SP's are specified by the P14 transgene in LckCre$^{+}$ P14$^{+}$ GSK-3αβ$^{fl/fl}$ and LckCre$^{-}$ P14$^{+}$ GSK-3αβ$^{fl/fl}$ mice while P14 transgene is undetectable in LckCre$^{+}$ P14$^{-}$ GSK-3αβ$^{fl/fl}$ CD3$^{+}$CD24$^{-}$CD8 SP cells.

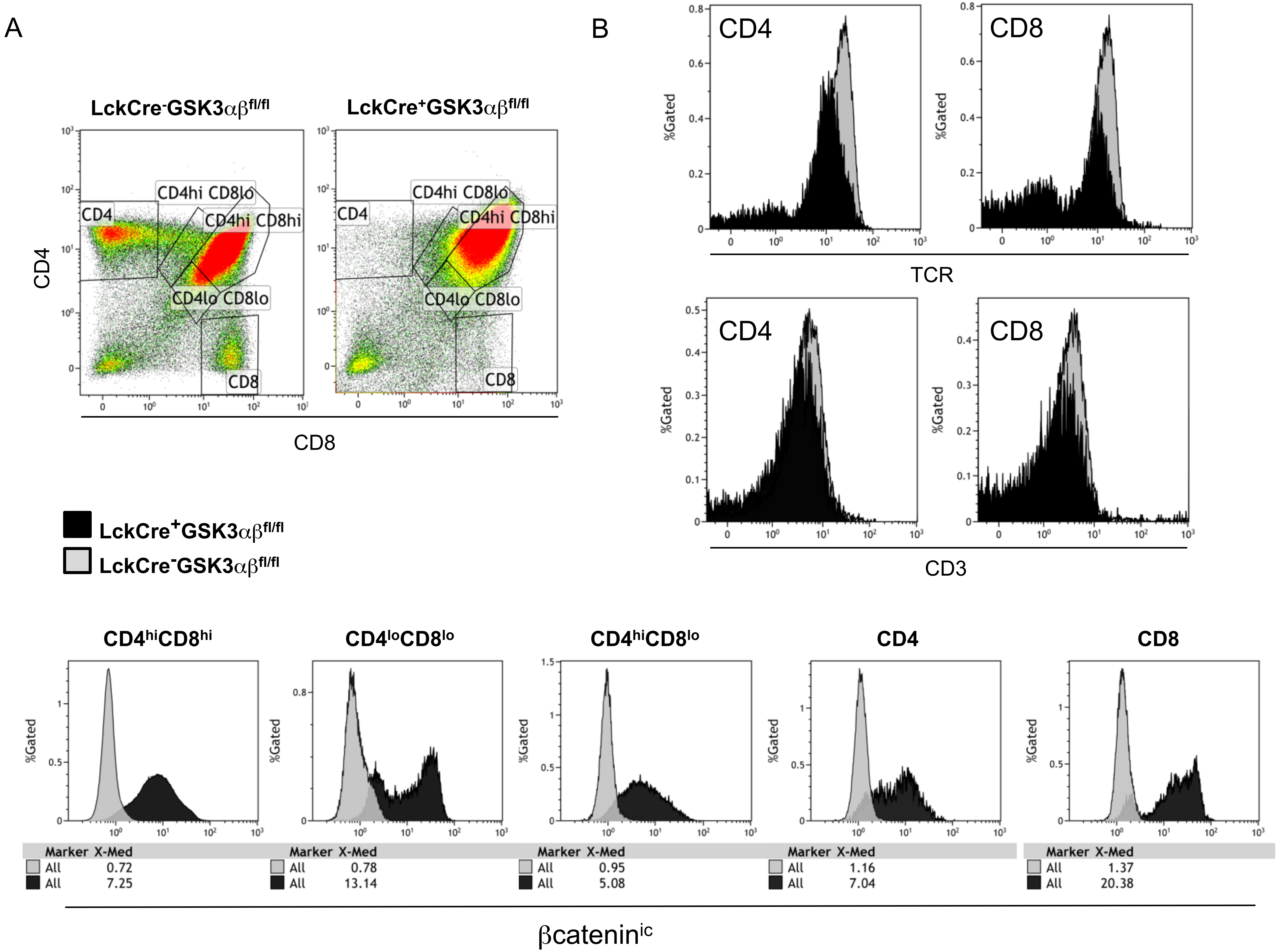


**Suppl. Figure 3.** A. Intracellular β-catenin expression expressed as median fluorescent intensity within gated thymocyte sub-populations. B. Splenic $CD4^{+}$ and $CD8^{+}$ SP populations from $LckCre^{+}$ GSK-3αβ$^{fl/fl}$ mice had comparable levels of TCR and CD3 expression to $LckCre^{-}$ GSK-3αβ$^{fl/fl}$ mice even though numbers were significantly reduced.

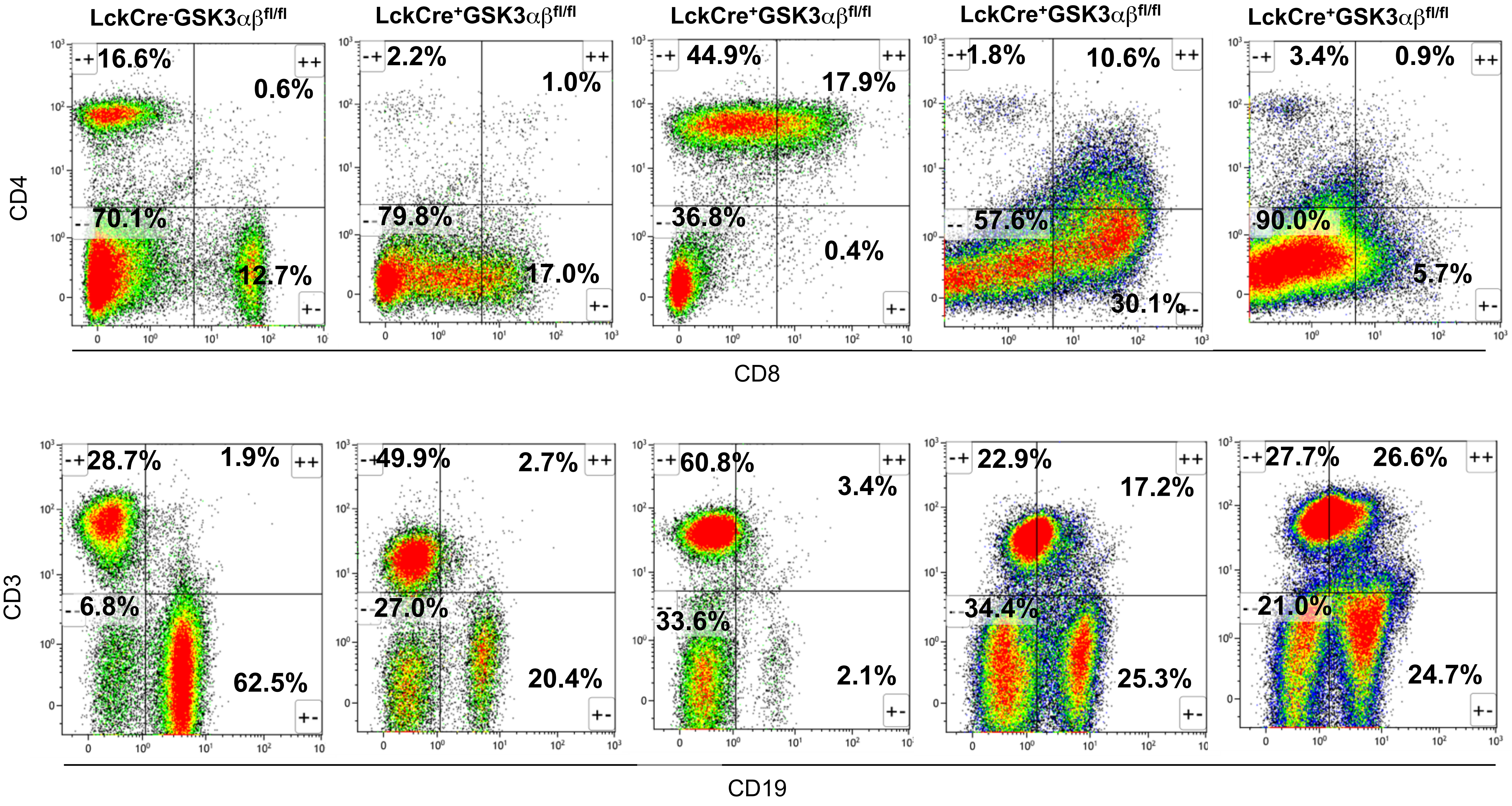


**Suppl. Figure 4.** FACS of 5 individual LckCre+ GSK-3αβ$^{fl/fl}$ lymphomas reveal heterogeneity as judged by CD3, CD4,CD8, and CD19 markers.
